# Interleukin-10-producing monocytes contribute to sex differences in pain resolution in mice and humans

**DOI:** 10.1101/2023.11.03.565129

**Authors:** Jaewon Sim, Elizabeth O’Guin, Karli Monahan, Chiho Sugimoto, Samuel A. McLean, Liz Albertorio-Sáez, Ying Zhao, Sophie Laumet, Andrew Dagenais, Matthew P. Bernard, Joseph K. Folger, Alfred J. Robison, Sarah D. Linnstaedt, Geoffroy Laumet

## Abstract

Pain is closely associated with the immune system, which exhibits sexual dimorphism. For these reasons, neuro-immune interactions are suggested to drive sex differences in pain pathophysiology. However, our understanding of peripheral neuro-immune interactions on sex differences in pain resolution remains limited. Here, we have shown, in both a mouse model of inflammatory pain and in humans following traumatic pain, that males had higher levels of interleukin (IL)-10 than females, which were correlated with faster pain resolution. Following injury, we identified monocytes (CD11b+ Ly6C+ Ly6G-F4/80^mid^) as the primary source of IL-10, with IL-10-producing monocytes being more abundant in males than females. In a mouse model, neutralizing IL-10 signaling through antibodies, genetically ablating IL-10R1 in sensory neurons, or depleting monocytes with clodronate all impaired the resolution of pain hypersensitivity in both sexes. Furthermore, manipulating androgen levels in mice reversed the sexual dimorphism of pain resolution and the levels of IL-10-producing monocytes. These results highlight a novel role for androgen-driven peripheral IL-10-producing monocytes in the sexual dimorphism of pain resolution. These findings add to the growing concept that immune cells play a critical role in resolving pain and preventing the transition into chronic pain.

**Graphical abstract:** 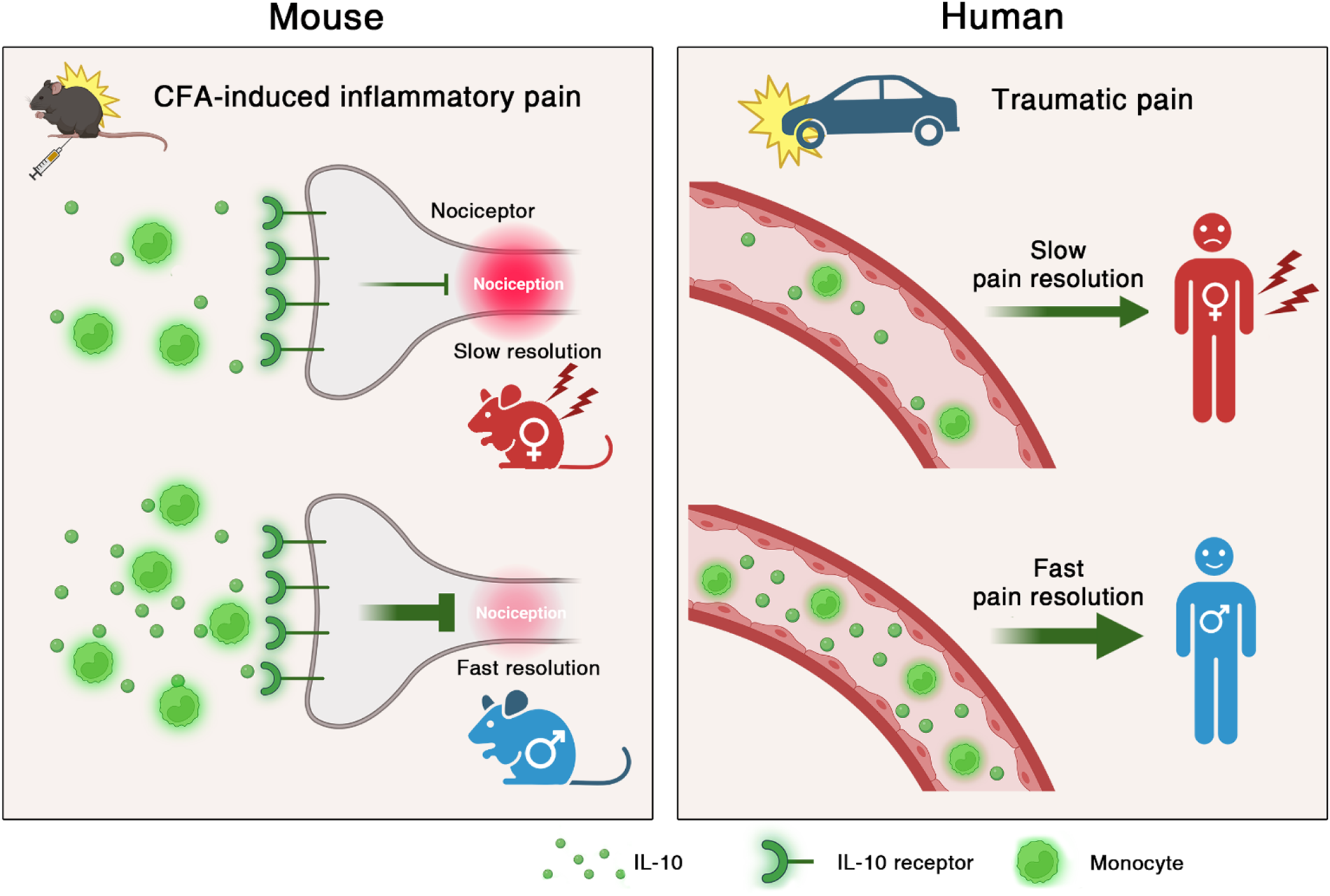

## Introduction

Chronic pain is a prevalent health issue that impacts more than 100 million individuals in the United States, costing $635 billion annually^1,2^. Women are disproportionately affected by chronic pain compared to men, as evidenced by epidemiological studies reporting female predominance in most chronic pain syndromes^3^. A higher risk of chronic pain in women is associated with a slower resolution of pain, causing females to experience a longer duration of pain^4–6^. To reduce the sexual inequality of pain-suffering, it is essential to gain a comprehensive understanding of the mechanisms that underlie this difference.

Given the tight association between the immune system and pain, sexual dimorphism in the immune response^7^ possibly contributes to sex differences in pain^8–10^. Cytokine receptors are widely expressed on nociceptors, facilitating direct communication between immune cells and sensory neurons to modulate pain^11^. For example, proinflammatory cytokines like TNFα, IL-1β, and IL-6 can sensitize nociceptors by increasing depolarizing currents, thereby inducing nociception^12–14^. In contrast, our previous research demonstrated that the anti-inflammatory cytokine IL-10 is crucial for reducing neuronal activity and pain hypersensitivity caused by cisplatin^15^. Although numerous studies have highlighted the importance of IL-10 in pain relief^16–18^, it is still unknown whether IL-10 directly contributes to the sex difference in pain resolution in rodents and humans. Furthermore, due to limitations in detecting IL-10 *in vivo* or *ex vivo*, the cellular source of IL-10 in pain conditions remains elusive.

We discovered that skin IL-10 plays a crucial role in resolving inflammatory pain induced by Complete Freund’s Adjuvant (CFA) in mice. In humans, higher levels of circulating IL-10 after traumatic injury predict faster pain resolution. Following injury, both in mice and humans, levels of IL-10 and monocytes were higher in males compared to females. We identified monocytes (CD11b+ Ly6G-F4/80^mid^ Ly6C+) as the primary source of IL-10 in the inflamed skin of mice by employing IL-10 GFP reporter mice. In humans, IL-10 levels are positively correlated only with the circulating level of monocytes, not with other cell types, suggesting that monocytes are the source of IL-10. In mice, IL-10 released by monocytes signals to IL-10R1 on advillin-expressing neurons. Blocking this neuro-immune communication impaired the resolution of pain hypersensitivity.

Previous studies revealed that sex hormones drive sex differences in immune response^7^ and pain^19–21^. Hence, we assessed the impact of sex hormones in mice on skin immune cells, IL-10 production, and the resolution of pain. Manipulations of androgen levels reversed the sexual dimorphism observed in pain resolution, infiltrated immune cells, and IL-10 production. Collectively, our findings suggest that androgen-driven peripheral IL-10-producing monocytes contribute to sexual dimorphism in pain resolution in mice and humans.

## Results

### Inflammation-driven skin monocyte infiltration is greater in males than females

CFA treatment strongly activates the immune system in the treated skin^22^. We observed an increase in the percentage of skin immune cells (CD45+) in the CFA-treated skin compared to the saline-treated skin (**Figure 1A, Supplemental Table 1**). To investigate the alterations in immune populations induced by CFA, we analyzed immune populations in the skin, identifying nine immune cell types: neutrophils (CD11b+ Ly6G+), monocytes (CD11b+ Ly6C+ Ly6G-F4/80^mid^), macrophages (CD11b+ F4/80^high^ Ly6C-Ly6G-), CD11b+ dendritic cells (CD11b+ CD11c+ F4/80-Ly6G-), eosinophils (CD11b+ SiglecF+ Ly6G-), CD11b-dendritic cells (CD11b-CD11c+ CD3-NK1.1-), granulocytes (CD11b-SSC^high^ CD3-NK1.1-), NK cells (CD11b-CD11c-CD3-NK1.1+), and T cells (CD11b-CD11c-CD3+ NK1.1+) (**Figure 1, B and C, Supplemental Table 1**). Saline injection did not significantly alter the skin immune populations from the naive skin, with macrophages, dendritic cells, and T cells being the predominant cell types in skin from both sexes (**Supplemental Figure 1A**). In contrast, CFA treatment completely reshaped the composition of immune cells in the skin, showing a remarkable increase in myeloid cells and lymphocytes (**Figure 1D, Supplemental Table 1**). The increase in monocytes was most significant, as further evidenced by the upregulation of monocyte marker genes as *Ccr2* and *Plac8* (**Supplemental Figure 1B,C**). Interestingly, we observed larger monocyte populations in males compared to females (p=0.0074) (**Figure 1E, Supplemental Table 1**). Collectively, these data demonstrate that there are more infiltrated monocytes in the inflamed skin of male mice than in females.

**Figure 1.**
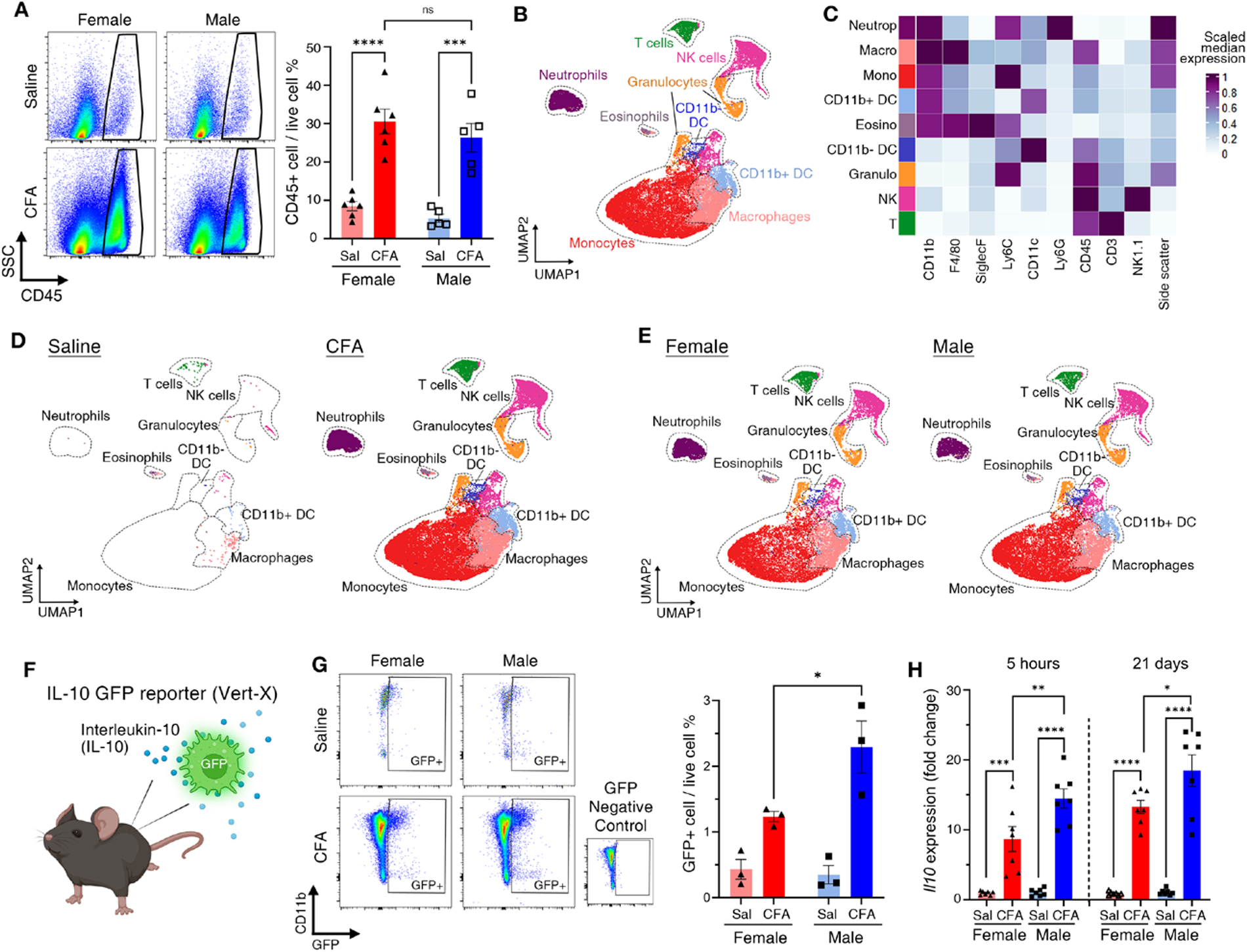
The immune populations in the skin differ between sexes under inflammation. (**A**) Flow cytometric detection of immune cells (CD45 positive) in the mouse plantar skin 7 days after treatment with saline or CFA (5 µL). Left, repre sentative flow cytometry plots presenting CD45 expression among live cells. Cells visualized on the dot plots are gated on FSC/SSC; Singlet; Live. Right, quantification of the percentage of immune cells in live cells in the skin. n=6 (Female), n=5 (Male). (**B** and **C**) Skin immune populations were identified with flow cytometry and high dimensional data analysis. (**B**) UMAP plot of skin immune cells treated with either saline or CFA. (**C**) Heatmap showing the relative expression of immune cell surface markers, with the color representing median expression each marker within each cluster. Dark purple, highest expression; White, lowest expression. (**D**) UMAP plot showing immune cell populations in saline (left) or CFA (right) treated skin. (**E**) UMAP plot showing immune cell populations in CFA-treated skin of female (left) or male (right). (**F**) Illustration of IL-10 GFP reporter mice. (**G**) Quantification of GFP+ (IL-10-producing) immune cells (CD45+) in the plantar skin of IL-10 eGFP mice, which was collected 7 days after saline or CFA treatment. Left, Flow cytometry plot showing a portion of GFP+ (IL-10-producing) immune cells among immune cells (pre-gated on CD45 positive liv e cells). Right, Quantification of the percentage of GFP+ immune cells in live cells. n=3 (Female), n=3 (Male). GFP+ cells were defined based on G FP negative control which was prepared from the CFA-treated skin of WT. (**H**) Relative *Il10* mRNA level in plantar skin 5 hours or 21 days after either saline or CFA treatment. n=7/group. Statistical analysis: two-way ANOVA, Tukey’s multiple comparisons (**A, G, H**). *p < 0.05, **p < 0.01. Error bar shows means and SEM (**A, G, H**). Abbreviation: Sal, saline; CFA, Complete Freund’s Adjuvant; Neutro, neutrophils; Macro, macrophages; Mono, monocytes; CD11b+ DC, CD11b positive dendritic cells; Eosino, eosinophils; CD11b-DC, CD11b negative dendritic cells; Granulo, granulocytes (mast cells and basophils); NK, NK cells; T, T cells. Two or three mice were pooled into one sample for flow cytometry experiments (**A** and **G**).

### IL-10 is higher in the skin of males than in females during inflammation

Skin IL-10 plays a crucial role in resolving inflammation and tissue repair after injury^23–25^. To identify IL-10-producing cells in the skin, we employed IL-10 GFP reporter mice (**Figure 1F**). The cells producing IL-10 in these mice co-express eGFP intracellularly, enabling us to detect the source of IL-10. We observed an increase in GFP+ immune cells in CFA-treated skin compared to saline-treated skin (**Figure 1G**). Importantly, CFA-treated male skin had a higher percentage of GFP+ immune cells than female skin (**Figure 1G**). Upregulation and sex differences in *Il10* gene expression in the CFA-treated skin were also observed at 5 hours and 21 days after CFA treatment (**Figure 1H**). CD45-negative cells did not express GFP, ruling out the possibility of IL-10 production from non-immune cells (**Supplemental Figure 1G**). *Il10* gene expression remained unaffected in the dorsal root ganglion (DRG), spinal cord (SC), and spleen (**Supplemental Figure 1D-F**). These results demonstrate that the upregulation of IL-10 was higher in males than females specifically in the CFA-treated skin.

### IL-10-mediated skin neuroimmune interactions resolve inflammatory pain

Mechanical pain hypersensitivity induced by CFA in mice persisted for 2-3 weeks. The resolution of pain hypersensitivity was significantly faster in males than in females, while the resolution of edema was similar in both sexes (**Figure 2A, Supplemental Figure 2A**). Interestingly, the percentage of GFP+ (IL-10-producing) cells and paw withdrawal threshold were positively correlated, showing that a higher number of IL-10-producing cells is associated with lower pain sensitivity (p<0.005) (**Figure 2B**). To test the casual role of IL-10 in the resolution of CFA-inflammatory pain, we inhibited skin IL-10 signaling by injecting neutralizing anti-IL-10 into the CFA-treated skin. Local IL-10 inhibition significantly delayed pain resolution in both sexes (**Figure 2C**). We observed the expression of IL-10 receptors in the sensory nerve terminals that innervate the plantar skin (**Figure 2D**). Based on these findings, we hypothesized that skin IL-10 may regulate pain resolution through IL-10 signaling in sensory neurons. To test the idea, we used *Avil^cre^:Il10ra^f/f^*(*Il10ra^DRG-KO^*) mice to specifically delete IL-10 receptors from sensory neurons (**Figure 2E, Supplemental Figure 2C**)^15^. Both sexes showed impaired pain resolution due to the absence of IL-10 signaling in advillin-expressing sensory neurons (**Figure 2F, Supplemental Figure 2B**). In wild-type (WT) mice, IL-10 receptor expression in sensory neurons did not differ between sexes and was unaffected by treatment (**Figure 2G, Supplemental Figure 2D**). Overall, these data demonstrate that inflammatory pain resolution is driven by neuroimmune interaction in the skin via immune cell-derived IL-10 acting on sensory neuron IL-10 receptors.

**Figure 2.**
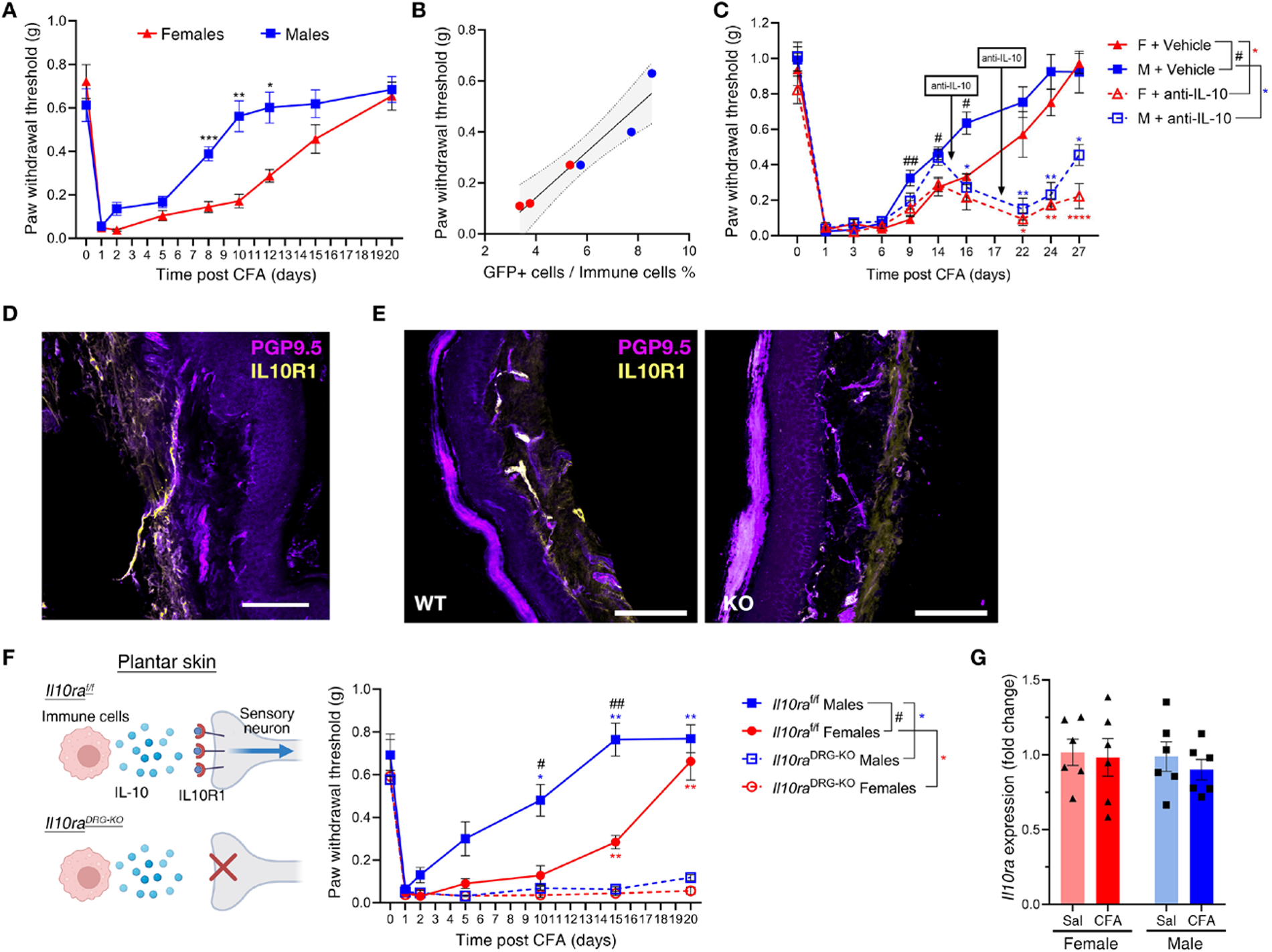
IL-10-mediated skin neuroimmune interactions resolve inflammatory pain. (**A**) Mechanical pain thresholds of WT females and males treated with CFA. n=5 (Female), n=7 (Male). (**B**) Correlation between percentage of GFP+ cells (IL-10 producing cells) in skin treated with CFA and paw withdrawal threshold measured on day 6, a day before skin collectio n from IL-10 GFP reporter mice. (Slope=0.09016, R^2^=0.9257, p=0.0021). Red dots: females, blue dots: males. n=3 (Female), n=3 (Male). 9 5% confidence interval bands of curves are shown as grey-filled areas. (**C**) Intraplantar injection of neutralizing anti-IL-10 on C57BL/6J (WT) females and males impeded the resolution of mechanical hypersensitivity in both sexes. Anti-IL-10 or vehicle was injected 14 and 17 days after CFA treatment. n=6/group. (**D**) Confocal images of plantar skin of C57BL/6J mouse, showing sensory nerve fiber expressing IL-10 rec eptor. Yellow (IL-10R1), Violet (PGP9.5). Scale bars, 100 μm. (**E**) Confocal images of plantar skin of *Avil^WT^:Il10ra^f/f^* (WT) mouse and *Avil^Cre^:I l10ra^f/f^* (KO) mouse. Yellow (IL-10R1), Violet (PGP9.5). Scale bars, 100 μm. (**F**) Mechanical pain thresholds of *Avil^WT^:Il10ra^f/f^* or *Avil^Cre^:Il10ra ^f/f^* females and males intraplantarly treated with CFA on day 0. Left, is an illustration describing the mouse model. Right, selective deletion o f IL-10 receptor in sensory neurons delays pain resolution. N=5 /group. *Il10ra^f/f^* (*Avil^WT^:Il10ra^f/f^*) and *Il10ra*^DRG-KO^(*Avil^Cre^:Il10ra^f/f^*). N=5 (each group). (**G**) Relative *Il10ra* mRNA level in lumbar dorsal root ganglion (DRG) 7 days after CFA treatment. N=6 / group. Statistical analysis: Ordinary two-way ANOVA, Tukey’s multiple comparisons (**G**), Repeated measures two-way ANOVA, Tukey’s multiple c omparisons (**A, C, F**), Simple linear regression (**B)**. *p or # < 0.05, **p or ## < 0.01, ***p < 0.005, ****p < 0.001, ns, not significant. Symbols represent means and error bars represent SEM (**A, C, F**). The error bar shows means and SEM (**G**). Abbreviation: Sal, saline; CFA, Compl ete Freund’s Adjuvant. Two or three mice were pooled into one sample for flow cytometry experiments.

### In humans, higher IL-10 levels in men are associated with faster pain resolution

Next, we evaluated the translational relevance of our preclinical findings. We examined human samples from the AURORA study^26^ involving participants with post-traumatic pain following motor vehicle accidents to assess sex differences in IL-10 levels and pain resolution. Circulating IL-10 levels and pain status were measured within 72 h after the traumatic injury, followed by additional pain severity assessments at 8 weeks and 3 months after the injury (**Figure 3A, Supplemental Table 3**). Men had significantly higher levels of circulating IL-10 than women after trauma (**Figure 3B**). Although men and women showed no differences in pain severity immediately after trauma, pain resolution was faster in men compared to women at 8 weeks and 3 months post-trauma (**Figure 3C**). We observed a negative correlation between the initial level of IL-10 and pain severity at 3 months, indicating that higher levels of IL-10 predict faster pain resolution (slope (β)= -0.455, p<0.05) (**Figure 3D, Supplemental Table 4**). This relationship was more pronounced in men (β= -0.666) than in women (β= -0.337) (**Figure 3E, Supplemental Table 5**). Overall, these results also showed that in humans, similarly to mice, men had higher levels of IL-10 correlating with faster pain recovery compared to women.

**Figure 3.**
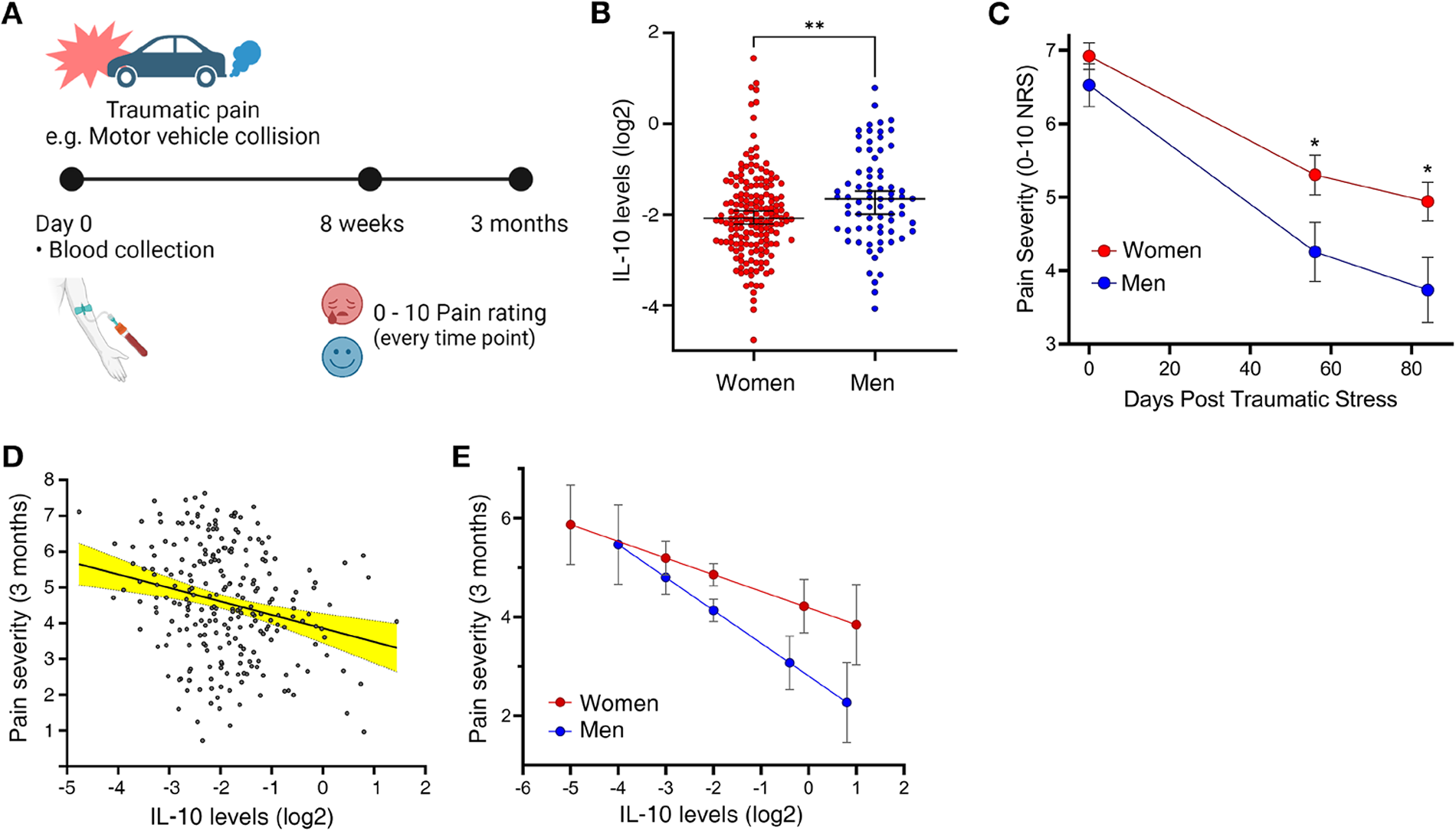
In humans, higher IL-10 levels in men are associated with faster pain resolution. (**A**) Experimental design of the AURORA study. (Women=172. Men=73). Day 0, when participants visited emergency departments within 72 hours after traumatic injury. (**B**) Plasma IL-10 levels in the immediate aftermath of traumatic injury. t=2.70, p=0.008. Error bars represent the median and 95% confidence interval. (**C**) Sex differences in the level of traumatic pain were compared on day 0 (t=-1.54, p=0.13), 8 weeks (t= -2. 37, p=0.019), and 3 months (t= -2.11, p=0.037) post-traumatic injury. (**D**) Relationship between circulating IL-10 levels (immediately after injury) and pain severity three months following injury. β= -0.455, p=0.031. 95% confidence interval bands of the curve are shown as yellow-filled areas. (**E**) The correlation between IL-10 levels and pain severity is higher in men compared to women. Women (β= -0.337, p=0.194) Men (β= – 0.666, p=0.118). Statistical analysis: Unpaired t-test (**B**), Repeated measures two-way ANOVA, Tukey’s multiple comparisons (**C**), linear mixed model regression (**D, E**). *p < 0.05. Symbols represent the median and error bars represent SEM (**C, E**). NRS, numeric pain rating scale.

### Monocytes are the primary source of IL-10 in mice during inflammatory pain

We next investigated which immune cells produce IL-10 in the inflamed skin of mice. In IL-10 GFP reporter mice, monocytes, macrophages, and T cells constituted the most abundant GFP+ (IL-10-producing) cell populations (**Figure 4A**). Monocytes were the most abundant GFP+ cells in both sexes, with males having significantly more GFP+ monocytes than females (males: 5.16%, females: 2.53%) (p<0.001) (**Figure 4B, Supplemental Table 2**). To determine which subset of monocytes produces IL-10, we categorized the monocytes/macrophages population as Ly6C^high^, Ly6C^mid^, and Ly6C^negative^ (**Supplemental Figure 3A**). Among them, about 70% were Ly6C^mid^ monocytes, indicating they are the primary cell type producing IL-10 in inflamed skin (**Supplemental Figure 3, B and C**). We also examined whether the percentage of GFP+ monocytes, macrophages, or T cells correlates with mechanical sensitivity. A higher percentage of GFP+ monocytes significantly correlated with less mechanical pain (p<0.01) (**Figure 4C**). No significant relationship was found for the other GFP-expressing cell types (**Supplemental Figure 3D,E**).

**Figure 4.**
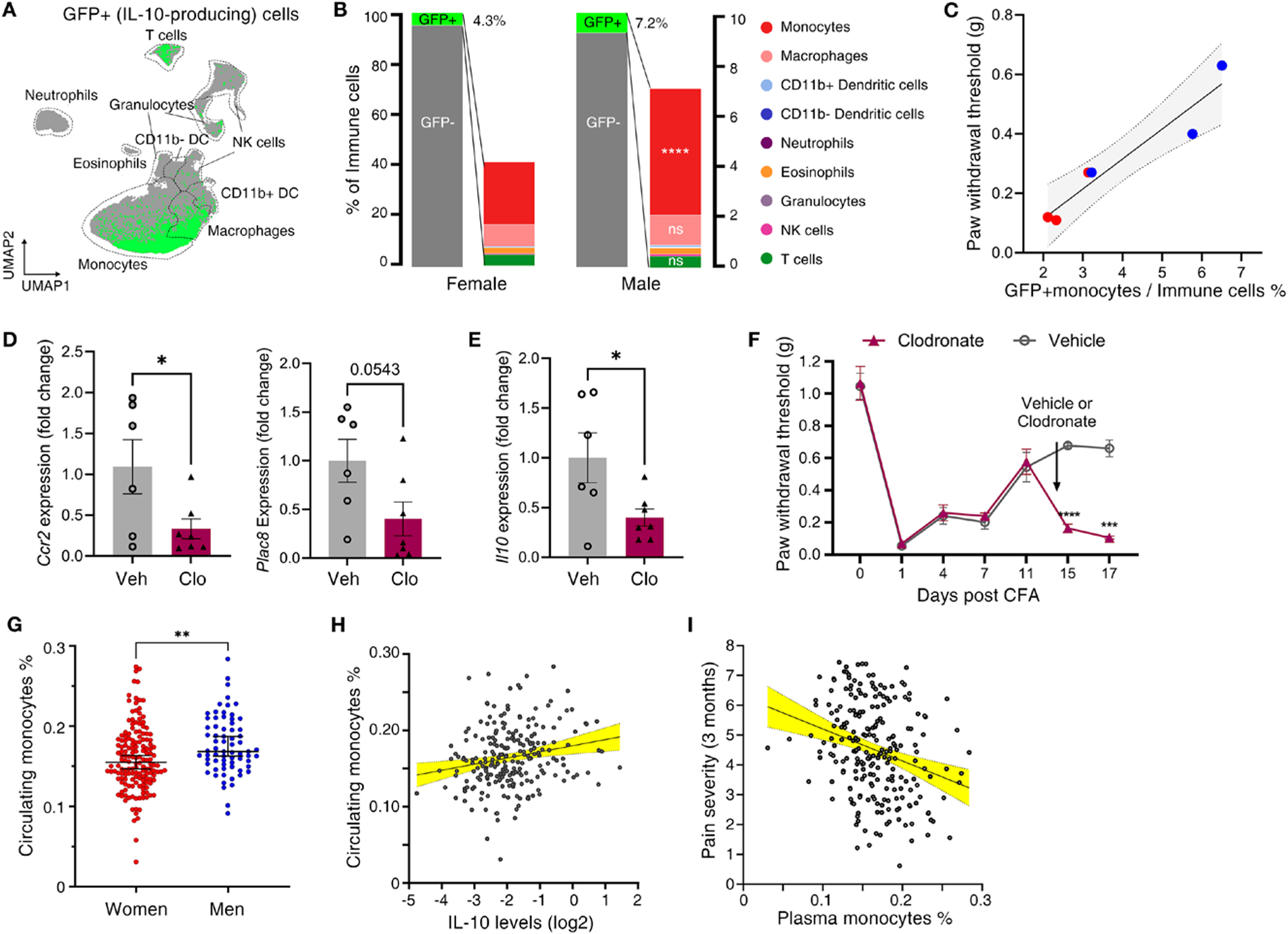
IL-10 is mainly produced from monocytes following painful injury in both mice and humans. (**A**) UMAP plot of skin immune cells from CFA-treated IL-10 GFP reporter mice. Green dots represent GFP+ (IL-10-producing) cells and gray dots represent GFP negative cells. (**B**) Bar graph showing the immune composition of GFP+ (IL-10-producing) cells in the CFA-treated skin. Mean valu es of three samples are only presented in this graph. SEM and significance are shown in **Supplemental Table 2**. The left y-axis applies to gray/gr een bars and the right Y-axis applies to compositions of GFP+ cells. Indication of significances represent statistical differences between sexes. n= 3 (Female), n=3 (Male). (**C**) Correlation between percentage of GFP+ monocytes (IL-10 producing monocytes) in CFA-treated skin and paw withd rawal threshold measured a day before skin collection. IL-10 GFP reporter mice were used, and flow cytometric analysis was conducted. (Slope= 0.1008, R^2^=0.9112, p=0.0031). 95% confidence interval bands of the curve are shown as grey-filled areas. Red dots: females, blue dots: males. n =3 (Female), n=3 (Male). (**D, E**) Relative mRNA levels of monocyte signature genes (*Csf1r, Plac8*) or *Il10* in plantar skin. Vehicle or clodronate lip osome was intraplantarly injected 14 days after CFA treatment. Skins were collected 3 days after clodronate treatment. n=6 (Vehicle), n=7 (Clodro nate). (**F**) Mechanical pain thresholds of C57BL/6J (WT) females and males intraplantarly treated with CFA and then clodronate liposome. N=7/gr oup. (**G**) CIBERSORT^100^– estimated percentage of human circulating monocytes in the immediate aftermath of traumatic injury exposure. t=3.14, p =0.002. The error bar represents the median and 95% confidence interval. N=245 (Women=172. Men=73). (**H**) Correlation between circulating IL-10 levels and percentage of monocytes immediately after the injury. rho=0.210, p=0.001. N=227. 95% confidence interval bands of the curve are shown as yellow-filled areas. (**I**) Correlation between circulating percentage of monocytes immediately after the injury and pain severity three mon ths following injury. β=-10.95, p=0.045. N=227. 95% confidence interval bands of the curve are shown as yellow-filled areas. Statistical analysis: Unpaired t-test (**D, E, G**), Repeated measures two-way ANOVA, Tukey’s multiple comparisons (**F**), simple linear regression (**C**), linear mixed models regression (**H, I**). *p < 0.05, **p < 0.01, ***p < 0.005, ****p < 0.001, ns, not significant. Symbols represent means and error bars represent SEM (**F**). Error bars represent means and SEM (**D, E**). Error bars represent median and 95% confidence intervals (**G**). Data show n in **G, H,** and **I** are from the human AURORA study. Abbreviation: DC, dendritic cell; Sal, Saline; Clo, clodronate liposome; Veh, Vehicle. Two or t hree mice were pooled into one sample for flow cytometry experiments.

Given that monocytes were the primary source of IL-10, we tested whether the depletion of monocytes can impact pain resolution. Skin monocytes were depleted by intraplantar injection of clodronate liposomes, and this was validated by the reduction of the expression of the monocyte markers *Ccr2* and *Plac8* (**Figure 4D**). Consequently, skin *Il10* gene expression was decreased after clodronate liposome treatment (**Figure 4E**). Clodronate-induced skin monocyte depletion impaired pain hypersensitivity resolution (**Figure 4F**), indicating that monocytes regulate pain resolution. Clodronate depletion did not change the expression of the neutrophil marker (*Ly6G*) and mast cell marker (*Cma1*) (**Supplemental Figure 3F,G**).

### Higher monocyte levels in humans are correlated with higher IL-10 and faster pain resolution

We again evaluated the translational value of our findings in the human AURORA study (**Figure 3A**). Comparably to mice, men had a higher percentage of circulating monocytes (estimated via RNA-seq) than women (**Figure 4G**). Additionally, we found a positive correlation between the percentage of monocytes and the level of circulating IL-10 (rho=0.210, p=0.001) (**Figure 4H**), while no correlation was found in other cell types such as T cells and dendritic cells (**Supplemental Figure 3I-K**). These results suggest that monocytes were the main producers of IL-10 (**Figure 4H**). The higher percentage of monocytes was also associated with less pain severity at 3 months, thereby accelerating pain recovery (β= -10.95, p=0.045) (**Figure 4I, Supplemental Table 6**). Further, we found, using formal mediation modeling, that IL-10 at least partially mediates the relationship between monocyte levels and pain (a=4.05, p=0.021; c’=-9.46, p=0.086) (**Supplemental Figure 3H**). Such relationships were not found in T cells (**Supplemental Figure 3L**). These results suggest that one potential mechanism through which monocytes contribute to pain resolution, in humans, might be their influence on IL-10 levels.

In summary, peripheral monocytes are plausibly the source of IL-10 and underlie sex differences in pain resolution in both mice and humans.

### Androgens regulate skin immunity and pain hypersensitivity resolution

We finally determined whether sex hormones mediate the sex differences observed in the resolution of pain, infiltration of monocytes, and IL-10 production. We gonadectomized mice to ablate sex hormones and supplemented ovariectomized (OVX) females with a placebo or dihydrotestosterone (DHT), a non-aromatizable metabolite of testosterone that activates the androgen receptor (**Figure 5A, Supplemental Figure 4A**). Body weight changes confirm DHT’s effects in females (**Supplemental Figure 4B**). Following CFA injection, pain resolution was not different between ovariectomized females treated with placebo and sham females (**Figure 5B**), indicating that estrogens are unlikely to play a role in sex differences in pain resolution. Meanwhile, OVX+DHT females exhibited faster resolution of mechanical hypersensitivity compared to other females (**Figure 5B, Supplemental Figure 4C**), while orchiectomized (ORX) males showed slower pain resolution than sham males (**Figure 5C, Supplemental Figure 4D**). This finding indicates that androgen hormones promote pain resolution.

**Figure 5.**
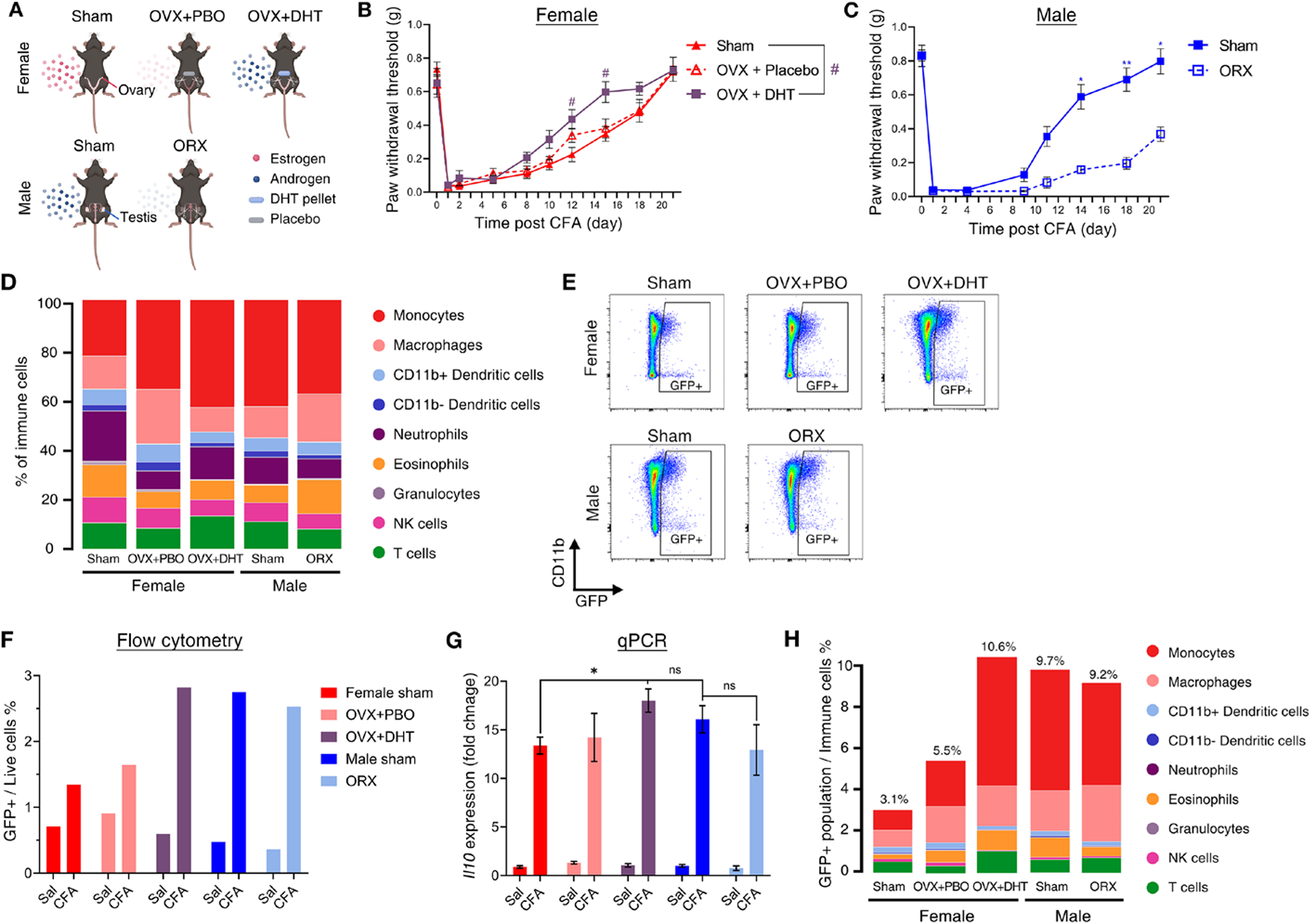
Sex hormone regulates skin immunity and pain hypersensitivity resolution. (**A**) Brief description on manipulating sex hormone levels in mice (C57BL/6J (WT) and IL-10 GFP reporter). (**B, C**) Mechanical pain thresholds of C57BL/6J (WT) females or males treated with CFA on day 0. Females; Sham (n=8), OVX + Placebo (n=8), OVX + DHT (n=8). Males; Sham (n=5), ORX (n=5). (**D**) Bar graph showing the immune cell populations in the CFA-treated skin of sex hormone-manipulated mice. (**E**) GFP+ (IL-10-producing) immune cells in the CFA-treated skin of sex hormone manipulated IL-10 eGFP mice. Flow cytometry dot plots showing a portio n of GFP (IL-10-producing) immune cells. Cells were pre-gated on CD45+ live cells. (**F**) Quantification of the percentage of GFP immune cells shown in **E**. Three CFA-treated plantar skins from individual animals were pooled into a single sample for each group. **(G)** Relative *Il10* mRNA level in the plantar skin of sex hormone-manipulated WT mice, 7 days after Saline or CFA treatment. Female: Sham (n=7), OVX + Placebo (n= 7), OVX + DHT (n=7). Male: Sham (n=5), ORX (n=5). (**H**) Bar graph depicting the immune composition of GFP+ cells (IL-10-producing) shown in **E** and **F**. Statistical analysis: Ordinary two-way ANOVA, Tukey’s multiple comparisons (**G**), Repeated measures two-way ANOVA, Tukey’s multiple comp arisons (**B, C**). *p or # < 0.05, **p < 0.01, ns, not significant. Symbols represent means and error bars represent SEM (**B, C**). Error bars show mean and SEM (**G**). Abbreviation: OVX, ovariectomy; PBO, placebo; DHT, dihydrotestosterone; ORX, orchiectomy; Sal, saline. Skin samples f rom three mice were pooled into one sample for flow cytometry experiments.

We assessed the impact of sex hormones on the skin immune response to CFA (**Figure 5D, Supplemental Table 7**). The percentage of skin monocytes was higher in OVX+DHT females than in other females and increased to levels comparable to those of sham males. We also evaluated the number of IL-10-producing cells using IL-10 GFP reporter mice. OVX females treated with a placebo were not different from sham females, indicating no role for estrogens. However, OVX+DHT females had the highest abundance of IL-10-producing cells in the inflamed skin among females, and the percentage of these cells was similar to that in sham males (**Figure 5E,F**). qPCR analysis confirmed the higher expression of *Il10* in OVX+DHT females (**Figure 5G**). Notably, the level of IL-10-expressing monocytes and the composition of IL-10-producing cells were similar between OVX+DHT females and sham males (**Figure 5H, Supplemental Table 8**). Manipulation of sex hormones did not affect *Il10* expression in the spinal cord (**Supplemental Figure 4E**). In conclusion, androgen promoted the production of IL-10 in the skin, leading to more rapid pain resolution.

Overall, our study revealed that sexual dimorphism in peripheral IL-10-producing monocytes underlies sex differences in pain resolution.

## Discussion

The precise biological mechanisms underlying the higher prevalence of persistent pain in females remain elusive. Our study discovered that the resolution of inflammatory pain requires the interaction of monocyte-derived IL-10 with IL-10R1 on advillin-expressing sensory neurons. Strikingly, we found that in both mouse and human males, the resolution of pain is faster and associated with more abundant IL-10 and monocytes. We propose a novel cross-species neuro-immune interaction contributing to sex differences in pain.

While neuro-immune interactions have been identified as significant contributors to chronic pain development^27,28^, an emerging body of studies has shown a critical role for the immune system in the resolution of pain^29–34^. Even in humans, activation of the immune system is necessary to resolve pain^35^. This supports the idea that immunosuppressive drugs are not the best response for treating chronic pain^36^. Consistently, we showed that immune cells, specifically monocytes, are essential for pain resolution, and depleting these cells with clodronate impaired the resolution of pain.

The analgesic role of IL-10 is well-established in preclinical studies^16,29,37,38^. Moreover, in humans, higher levels of circulating IL-10 are associated with reduced pain^39–43^. However, a mechanistic explanation widely applicable from animal models to humans has not yet been demonstrated. We found that IL-10 underlies the sexual dimorphism of pain duration in mice and humans. In our models, the major cellular source of IL-10 was monocytes, a cell type that was more prevalent in males of both species, consistent with previous literature showing male predominance in monocyte numbers in mice^44,45^ and humans^46,47^. Our findings show an overarching mechanism of pain resolution and sex differences, suggesting enhancing IL-10-producing monocytes as novel therapeutic strategies for the prevention and treatment of pain.

Our preclinical data indicate that IL-10 acts on DRG neurons to resolve inflammatory pain. In line with our findings, previous reports showed the critical role of IL-10 signaling in somatosensory neurons for alleviating pain^48,49^. While further evidence is needed in this model, it is plausible that IL-10 alleviates pain by reducing the spontaneous hyperactivity of DRG neurons^15^ or inhibiting the upregulation of voltage-gated sodium channels in these neurons^50^. Remarkably, IL-10 receptors are predominantly expressed in nonpeptidergic nociceptive neurons^51^, which are known to contribute to inflammatory mechanical hypersensitivity^52^. Therefore, it is likely that IL-10 resolves inflammatory pain by reducing the excitability of these nociceptors.

Due to the broadly expressed IL-10 receptors in immune cells, IL-10 is a potent immunosuppressive cytokine that might also relieve pain by dampening inflammation^18,53–55^. Our findings, however, shed more light on the direct action of IL-10 on sensory neurons. This is supported by the result that the ablation of IL-10R1 on advillin-positive sensory neurons was sufficient to prevent pain recovery. The unaffected edema size in this model also suggests that the inflammation status itself remained unchanged, while pain resolution greatly differed between WT and IL-10R-ablated mice.

Determining the cellular source of IL-10 remains a technical challenge due to the lack of adequate antibodies. We combined high-dimensional spectral flow cytometry with IL-10 GFP reporter mice to overcome this challenge and identified several immune cell types producing IL-10 in the inflamed paw: monocytes, macrophages, and T cells. Surprisingly, more than half of the IL-10-producing cells were classical Ly6C+ monocytes. The role of monocytes in pain is understudied, and they are mostly seen as macrophage precursors. However, it is now clear that monocytes contribute to inflammatory and physiological functions on their own^56,57^. Here, we showed the essential role of monocytes in pain resolution and sex differences in pain. Monocytes exhibit different phenotypes; high expression of Ly6C is generally considered proinflammatory^22,58,59^, while Ly6C^low^ or Ly6C^mid^ monocytes in tissue, often have an anti-inflammatory phenotype^58,58,60^. We found that Ly6C^mid^ monocytes were the major IL-10 producer, not Ly6C^high^ monocytes. Therefore, our results consistently showed that Ly6C^mid^ monocytes are protective, mediating IL-10-driven-pain resolution.

The importance of monocyte/macrophage populations in regulating inflammatory pain has been identified in other studies based only on a single sex^29,37^. Similar to our findings, the depletion of monocyte/macrophage cells in either zymosan-or carrageenan-induced inflammatory pain prolonged the duration of pain. However, a comparison of this mechanism between sexes has not been previously conducted. Our study is among the first to show that a higher number of IL-10-producing monocytes in males than in females underlies sexual dimorphism in pain resolution. This is consistent with previous literature that has shown that females experience greater inflammatory pain than males^34,55,61–64^. We specifically highlighted that males exhibit a significantly faster resolution compared to females. Similarly, a longer duration of pain in females than in males is broadly observed in both humans^4–6^ and animal models^61,65,66^. Our results imply that sex differences in monocytes contribute to the faster pain resolution in males.

The unequal number of IL-10-producing monocytes could possibly be attributed to sex hormones. Monocytes possess androgen^67,68^ and estrogen receptors^69^, enabling the modulation of these cells through sex hormones. In humans, the administration of testosterone increases the number of circulating monocytes^70^, while estrogen has the opposite effect^71,72^. Testosterone also influences the phenotype of monocytes, shifting them towards an anti-inflammatory phenotype^68,73,74^, thereby increasing IL-10 production^75–77^. Our study revealed that the administration of testosterone in females could increase the number of IL-10-producing monocytes, consequently facilitating faster pain resolution. This finding aligns with previous literature that has demonstrated the protective role of testosterone in inflammatory pain in mice^78,79^ and human studies which demonstrated a negative correlation between testosterone levels and the severity of pain^80,81^. However, we also observed a significantly delayed recovery in orchiectomized male mice, even though their IL-10 levels were marginally decreased compared to sham males. This could be attributed to the presence of androgen receptors on neurons^82–85^. While the role of androgen signaling in these neurons in reducing pain has not been extensively studied, it is plausible that testosterone exerts antinociceptive effects by modulating the nervous system^85,86^, in addition to the immune system. Interestingly, removing estrogen from female mice did not significantly alter pain status or IL-10 levels. The role of estrogen in pain regulation, as well as IL-10 production, remains complex and enigmatic. While many pieces of evidence consistently support testosterone’s protective role in pain relief^78–81^, contradictory results have emerged concerning the action of estrogen in pain modulation in both rodents^87–92^ and humans^20,93,94^. Reports on the impact of estrogen on IL-10 production are also similarly perplexing and contradictory^71,95–99^.

The first limitation of our study is that pain assessments between animals and humans are qualitatively different. While the von Frey test applied to mice measures evoked mechanical hypersensitivity, the numeric rating scale conducted on humans relies on self-reporting of pain intensity experienced in their daily lives. Additionally, we examined humans suffering from pain due to traumatic injuries, while local inflammatory pain was induced in mice. The second limitation is that cell types in human samples were estimated by CIBERSORT, based on RNA expression patterns. While this method is well-validated^100^, it is not as accurate as direct measurements of cell types by flow cytometry. Therefore, our inability to directly identify the cellular source of IL-10 in humans deters us from making a definitive conclusion that IL-10 is mainly produced from monocytes in humans. We found a positive correlation between IL-10 levels and monocytes but not with T cells, CD4+T cells, or dendritic cells in the large sample size (n=245). The absence of a positive correlation, especially between IL-10 levels and CD4 T cells, which are generally known as a major source of IL-10, contrasts with the remarkably strong correlation found in monocytes. Although correlation is not a direct measure of the source of IL-10, the striking association between monocytes and IL-10 levels supports the possibility that the production of IL-10 from monocytes underlies the statistically positive correlation. These differences between preclinical and clinical studies may hinder an exact comparison between the two. However, we consistently identified sex differences in pain, as well as an association of pain with IL-10 and monocytes with different pain assessments in two distinct organisms with different pain conditions. This finding strengthens the idea that the mechanism we proposed here is broadly applicable in various pain models and conditions.

Overall, our study highlights the role of IL-10 and IL-10-producing monocytes in mediating neuroimmune interaction in the skin to relieve pain. It also provides valuable resources on skin immunity during CFA-induced inflammation. Our study features skin immune cells as potential targets and IL-10 as a key molecule for pain treatment. Given the high accessibility of the skin for clinical treatment and diagnosis compared to the nervous system, uncovering the pain resolution mechanisms in the skin could facilitate the development of clinical treatments and broaden their application. Importantly, our study reveals a remarkable consistency between the results of preclinical and clinical studies. We emphasize the critical role of IL-10-producing monocytes in pain resolution and highlight the importance of utilizing both sexes and incorporating human data to enhance the translational relevance of preclinical findings.

## Methods

Full methods are available in the Supplemental Methods

### AURORA study: Human participants

All human participants included in the current analyses were participants of the AURORA study, a large longitudinal study of adverse neuropsychiatric outcome development following trauma exposure^26^. Enrollment for this study took place between September 2017 and June 2021. Participant enrollment occurred at 23 urban emergency departments across the US, in the immediate aftermath of traumatic collision exposure, when the individual presented for care. All 245 participants included in the current analyses provided informed consent, completed baseline assessments in the emergency department, and provided blood samples for plasma and RNA collection. This is a diverse cohort including women and men from both black and white individuals. The population includes only these two self-identified racial/ethnic groups. **Supplemental Table 3** describes the characteristics of the studied population.

### Human pain assessment and outcome definition

Overall musculoskeletal pain severity was assessed in the immediate aftermath of traumatic collision exposure and at eight weeks and three months following enrollment by asking participants to rate their pain severity in the past week via a 0-10 numeric rating scale (NRS: 0 (no pain) to 10 (maximum possible pain))^101^.

### MSD Analysis of Cytokines

Human plasma cytokines (IL-6, IL-8, IL-10, TNFα, and IFNγ) were measured in multiplex using Meso Scale Discovery’s V-PLEX Proinflammatory Panel 1 Human chemiluminescence-based assay (MSD, Gaithersburg, MD, USA, catalog # K15049D-1).

### Human RNA collection, sequencing, and CIBERSORT

To estimate the proportions of immune cell types in each participant, we used RNA sequencing data from the AURORA cohort and a bioinformatic method of identifying cell types based on RNA expression profiles called CIBERSORT^100^.

### Animals

Mice were obtained from Jackson Laboratory (Bar Harbor, USA) and bred at Michigan State University animal facility. The following mouse strains were used: C57BL/6J (WT, JAX 000664), B6(Cg)-Il10^tm1.1Karp/J^ (Il10^GFP^, JAX 014530)^102^, B6.129P2-Avil^tm2(cre)Fawa^/J (Advillin^Cre^, JAX 032536)^103^, and B6(SJL)-Il10ra^tm1.1Tlg^/J (IL-10Rα^flox^, JAX 028146)^104^. Mice had *ad libitum* access to food and water and were housed in a light-dark cycle (12 h, each) under the condition of controlled humidity and temperature. Mice at 15 to 25 weeks of age were used in every experiment. Avil^Cre^:*Il10ra ^f/f^* mice were generated to genetically remove IL-10 receptors from the sensory neurons as previously described^15^.

### Animal model

To induce inflammatory pain, female and male mice received 5 µl of Complete Freund’s adjuvant (CFA, 1 mg/ml, F5881, Sigma-Aldrich, St. Louis, MO) in the plantar skin of hind paw, under anesthesia (isoflurane 2.5%). Control mice received the same volume of saline under anesthesia. Mechanical pain sensitivity threshold was measured using von Frey filaments as previously described^105–107^.

### Single cell suspension & flow cytometry

Mouse skin immune cells were isolated as previously described using enzymatic and mechanical digestion^22,108^. Cells were stained with flow cytometry antibodies (**Supplemental Table 9**) in flow cytometry staining buffer (PBS, 1% BSA) and brilliant staining buffer (563794, BD, Franklin Lakes, NJ). Cells were analyzed using a 5-laser (16UV-16V-14B-10YG-8R) Cytek® Aurora – Spectral Flow Cytometry located in the MSU Flow Cytometry Core Facility (Cytek Biosciences, Fremont, CA), and the data were unmixed with SpectroFlo® software and analyzed with FlowJo™ v10.8 Software (BD Life Sciences).

### High dimensional data analysis of flow cytometry data

Flow cytometry data were analyzed by using previously established workflow using R(v.4.2.0) and RStudio(2022.02.2+485)^109,110^.

### Illustration

Illustrations in Figure 1F, 2F, 3A, 5A, Supplemental Figure 1D, 1E, and 5A were created with BioRender.com.

### Statistics

Mouse flow cytometry, behavioral, qPCR, and ELISA data are presented as mean ± standard errors. Groups were considered statistically different when p<0.05 following appropriate statistical correction based on experimental design. The statistical test used is described in the figure legends. Statistical differences are illustrated as follows *p < 0.05, **p < 0.01, ***p < 0.005, ****p < 0.001, ns, not significant. Test and statistical differences and graphics were made with GraphPad Prism 9.

Human data are presented as median ± 95% confidence intervals. Sociodemographic characteristics of the human study were summarized using standard descriptive statistics. Human circulating IL-10 levels were determined to be non-normally distributed via the Shapiro-Wilk test; therefore, IL-10 levels were log-transformed by taking the natural log of raw IL-10 values. Linear mixed models were used to assess the relationship between peritraumatic IL-10 levels and chronic posttraumatic pain levels using the lmer function from the lme4 package (v. 1.1.23)^111^ in R (v. 3.6.2)^110^. Based on the results of previous studies (e.g.^112,113^), participant age, self-identified race/ethnicity, area deprivation index, and baseline pain levels were included in the models as covariates.

### Study approval

Animal: All preclinical experiments were approved by the Institutional Animal Care and Use Committee at Michigan State University and in accordance with the guidelines from the National Health Institute (NIH). Human: All participants provided written informed consent prior to enrollment and were compensated for their study participation^26^. The project was approved by each participating Institutional Review Board.

### Data availability

Preclinical and clinical raw data will be made available upon reasonable request. In addition, all AURORA study data will be available via the NIMH Data Archives.

## Author contributions

J.S. performed high-dimensional flow cytometry and analyzed the data. G.L., J.S., and S.L. performed behavioral assessments. M.P.B. provided expertise in flow cytometry. E.O., K.M., and A.D. performed mouse tissue collection and molecular experiments (RNA isolation/ qPCR/ ELISA). J.K.F. and S.L. bred mouse lines. C.S. and A.J.R. performed the gonadectomy. J.S. and S.L. performed intraplantar injection. J.S. and G.L. performed immunohistochemistry. J.S. made figures. J.S. and G.L. analyzed the preclinical data. G.L. and S.D.L. supervised the study. S.A.M. was the lead investigator on the AURORA study and was responsible for participant enrollment and follow-up. S.D.L. led the collection, processing, and analysis of biological specimens and associated biological data from AURORA study participants. L.A.S. isolated RNA from participant blood samples and measured cytokines in participant plasma. Y.Z. and L.A.S. performed data cleaning, statistical analyses, and visualization of human cohort data. J.S. and G.L. wrote manuscript drafts, all authors contributed to the discussion and edited the final version of the manuscript.

## Acknowledgements

This study was supported by the US National Institute of Health (NIH) R01 NS121259 (GL), U01 MH110925 (SAM), K01 AR071504 (SDL), R01 NS118563 (SDL and SAM), R01 MH111604 (AJR), The Rita Allen Foundation (GL and SDL), MSU Neuroscience DRF (GL), and the Department of Defense CP#220093 (GL).

The content is solely the responsibility of the authors and does not necessarily represent the views of these funding agencies. The authors would like to thank the participants for taking part in this study. The authors would like to acknowledge the University of North Carolina BioSpecimen Facility for the storage, accessioning, and disbursement of biological samples.

We thank Dr. Stephanie Puig (Boston University) for expert advice on skin immunostaining.

## Supplementary Materials

### Supplementary Figures

**Supplemental Figure 1.**
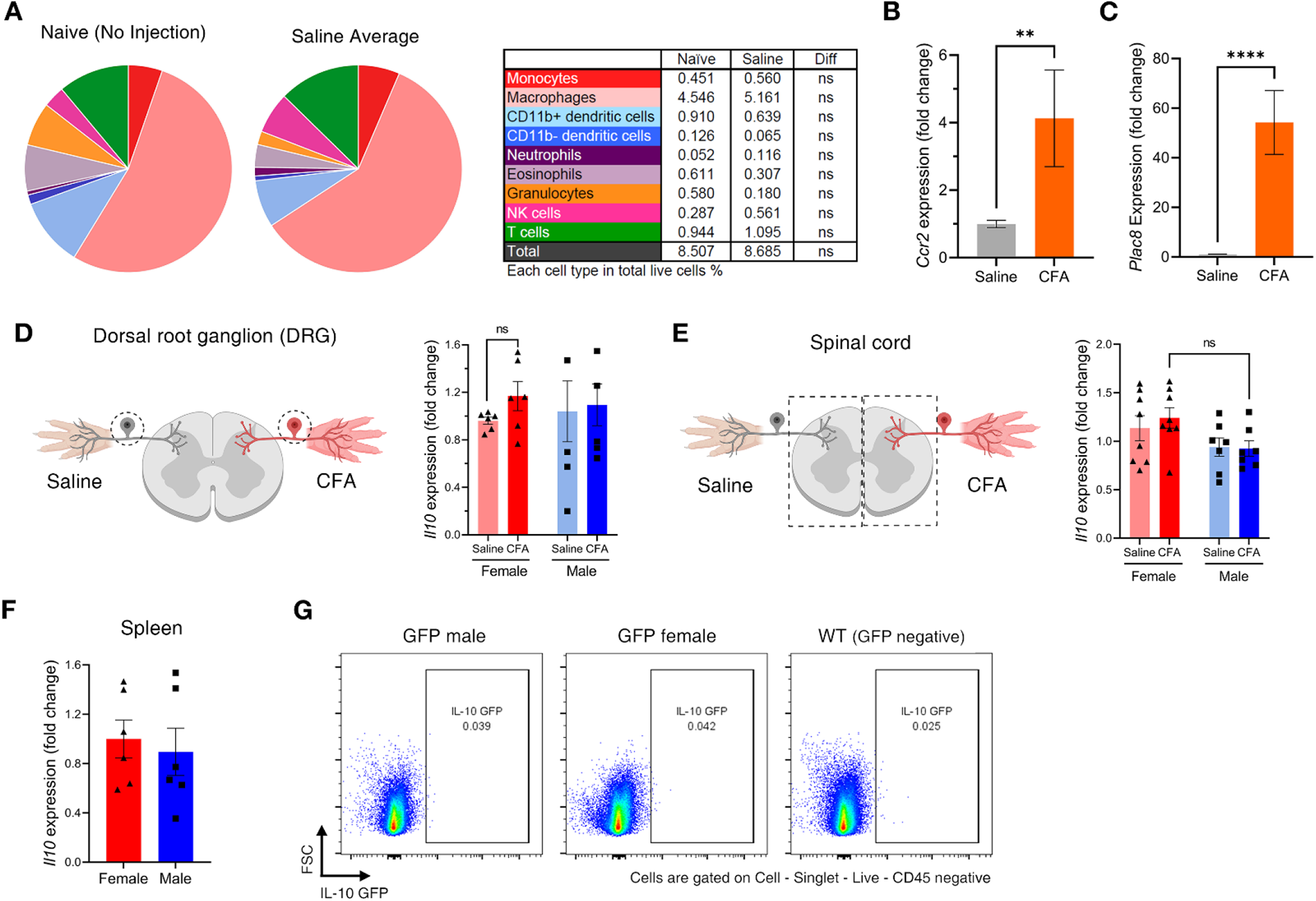
Expression of IL-10 in other tissues. **(A)** Pie charts showing the proportion of skin immune cells in the plantar skin without any treatment (naive) or 7 days after saline injection. The table shows the percentage of each cell type and a statistical comparison between naive and saline-treated skin. (**B** and **C**) Relative mRNA expression of monocyte marker genes *Ccr2 and Plac8* levels in saline or CFA-treated plantar skin 17 days after the treatment. Saline (n=14), CFA (n=6). (**D**) Relative *Il10* mRNA level in lumbar dorsal root ganglion (DRG) 7 days after CFA treatment. n=6/group. (**E**) Relative *Il10* mRNA level in the lumbar spinal cord 1 day after CFA treatment. n=8 (female), n=7 (male). (**F**) Relative *Il10* mRNA level in spleen 7 days after CFA treatment. n=6/group. (**G**) Flow cytometry dot plot showing a portion of IL-10-producing cells (GFP cells) among non-immune cells (gated on live CD45 negative cells). Statistical analysis: two-way ANOVA, Tukey’s multiple comparison (**A, D, and E**). unpaired t-test (**B, C**, and **F**). *p < 0.05, **p < 0.01, ns, not significant. Error bar shows means and SEM (**B-F**).

**Supplemental Figure 2.**
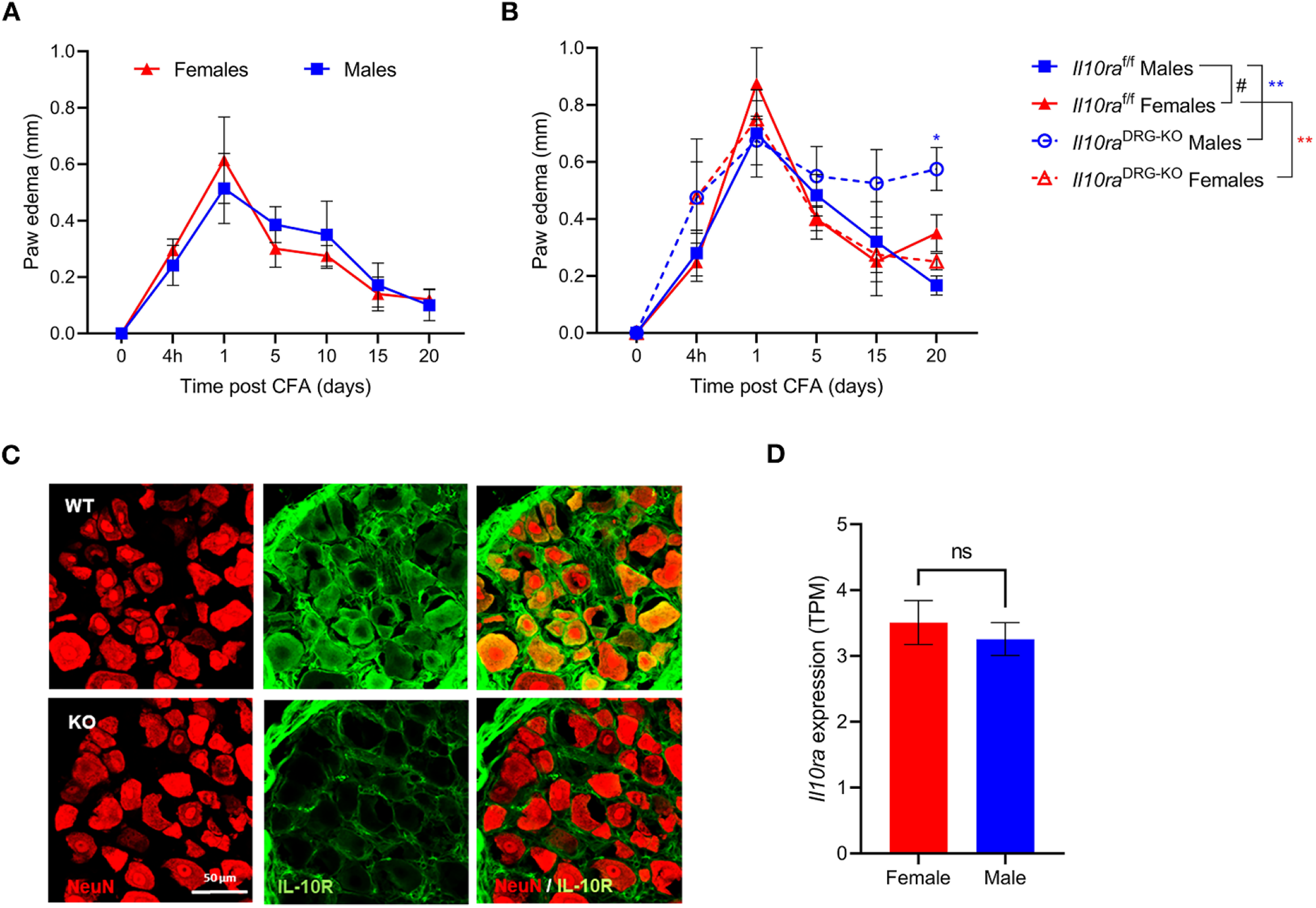
Expression of IL-10R1 in dorsal root ganglion. The size of paw edema after CFA treatment in (**A**) C57BL/6J (WT) female (n=5) and male (n=7) and *Avil^WT^:Il10ra^f/f^* and *Avil^Cre^:Il10ra^f/f^*. *Il10ra^f/f^*, *Avil^WT^:Il10ra^f/f^*; *Il10ra*^DRG-KO^, *Avil^Cre^:Il10ra^f/f^*. N=5 (each group). (**C**) Confocal images of DRG of *Avil^WT^:Il10ra^f/f^* and *Avil^Cre^:Il10ra^f/f^*. Red, NeuN; Green, IL-10R1. WT, *Avil^WT^:Il10ra^f/f^* ; KO, *Avil^Cre^:Il10ra^f/f^* . Scale bar, 50 μm. (**D**) Published RNA-seq data confirms the absence of sex difference in *Il10ra* mRNA levels in mouse DRG. doi: 10.1097/j.pain.0000000000001866. TPM, Transcript per million. N=6 (each group). Statistical analysis: Repeated measures two-way ANOVA, Tukey’s multiple comparisons (**A, B**), unpaired t-test (**D**). ns, not significant. Symbols represent means and error bars represent SEM (**A, B**). Error bars show mean and SEM (**D**).

**Supplemental Figure 3.**
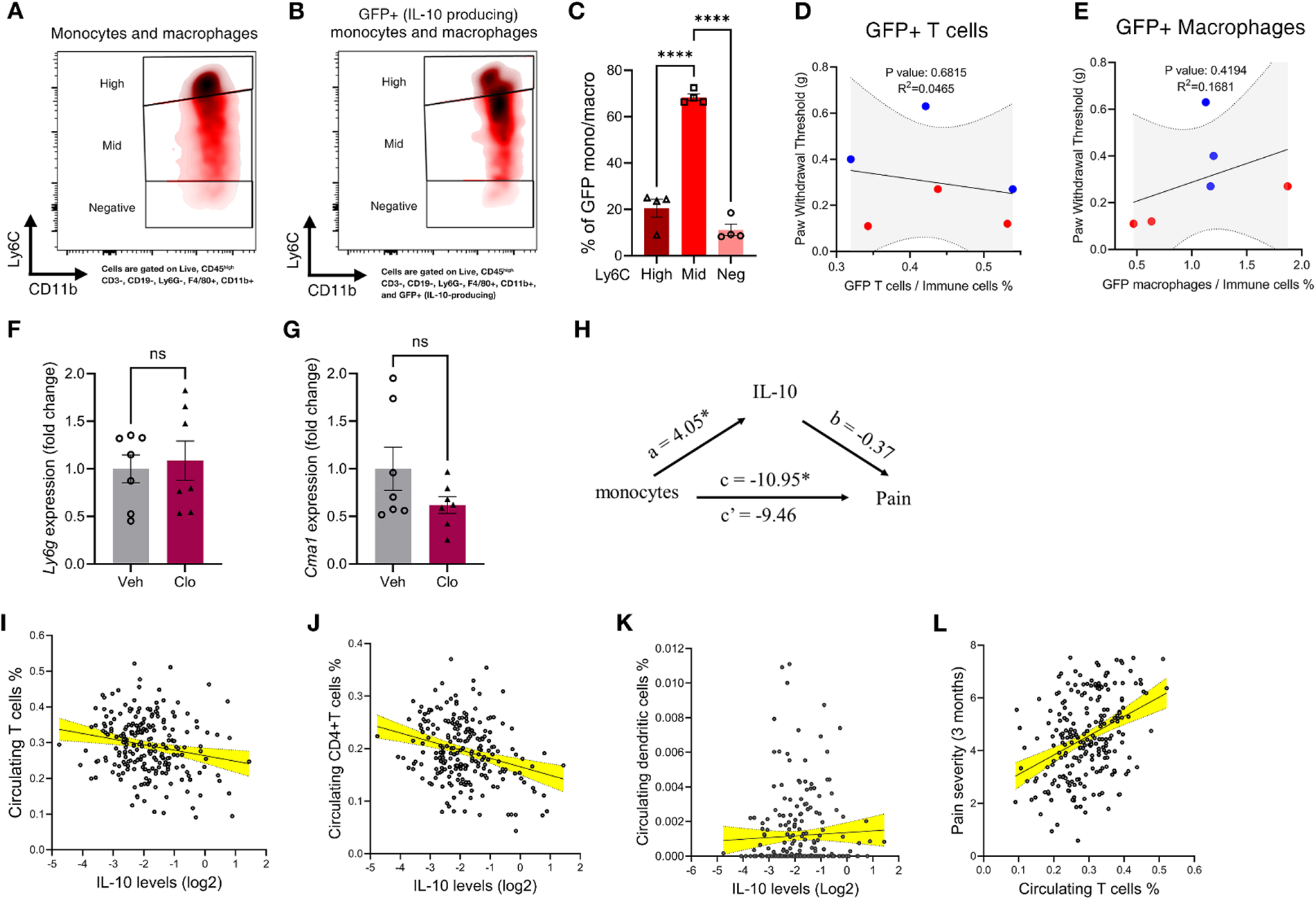
The majority of IL-10-producing monocytes were Ly6C^mid^ in mice and IL-10 levels do not correlate with dendritic cells and T cells in humans. (A) Flow cytometric density plots showing Ly6C intensity in monocytes/macrophages in CFA-treated plantar skin. Monocytes/macrophages are gated on live immune cells and CD11b+ Ly6G-CD11c-F4/80+. These populations were divided into three subtypes: Ly6C high, Ly6C mid, and Ly6C negative. (B) Similar to (**A**), but cells were gated on GFP+ (IL-10-producing) live immune cells and CD11b+ Ly6G-CD11c-F4/80+. (**C**) Percentages of three subtypes in GFP+ (IL-10-producing) monocytes/macrophages population. Skin samples from 4 mice were analyzed. (**D, E**) Correlation between percentage of GFP+ (IL-10-producing) T cells or macrophages in CFA-treated skin and paw withdrawal threshold measured on day 6, a day before skin collection from IL-10 GFP reporter mice. GFP T cells: Slope=1.032, R^2^=0.04652, p=0.6815. GFP macro: Slope=0.1611, R^2^=0.1681, p=0.4194. 95% confidence interval bands of curves are shown as grey-filled areas. Red dots: females, blue dots: males. N=6. (**F, G**) Relative mRNA levels of neutrophil (*Ly6g*) and mast cells (*Cma1*) signature genes in plantar skin. Vehicle or clodronate liposome was intraplantarly injected 14 days after CFA treatment. Skins were collected 3 days after clodronate treatment. n=6 (Vehicle), n=7 (Clodronate). Data shown in (**H-L**) are from the human AURORA study. (**H**) Statistical mediation analysis on the relationship between monocytes (intervention), IL-10 (mediator), and pain severity after three months (outcome). a, how the intervention changes the mediator; b, how the mediator impacts the outcome; c, how the intervention is related to the outcome in the presence of the mediator; c’, how the intervention is related to the outcome in the absence of the mediator^1^. a (T=1.742, p=0.021), b (T=1.69, p=0.09), c (T=5.438, p=0.045), c’ (T=5.484, p=0.086). (**I, J, K**) Correlation between circulating IL-10 levels and percentage of circulating T cell, CD4 T cell, and dendritic cells immediately after injury. N=227. (**I**) rho=-0.195, p=0.003, (**J**) rho=-0.236, p=0.0003, (**K**) rho=0.08, p=0.229. 95% confidence interval bands of the curve are shown as yellow-filled areas. (**L**) Relationship between circulating percentage of T cells (immediately after injury) and pain severity three months following injury. rho= 0.164, p=0.014. 95% confidence interval bands of the curve are shown as yellow-filled areas. Error bars represent mean and SEM (**C, F, G**). Statistical analysis: Ordinary one-way ANOVA, Tukey’s multiple comparisons (**C**), unpaired t-test (**F, G**), simple linear regression (**D, E**), linear mixed models regression (**I, J, K, L**). *p < 0.05, **p < 0.01, ***p < 0.005, ****p < 0.001, ns, not significant.

**Supplemental Figure 4.**
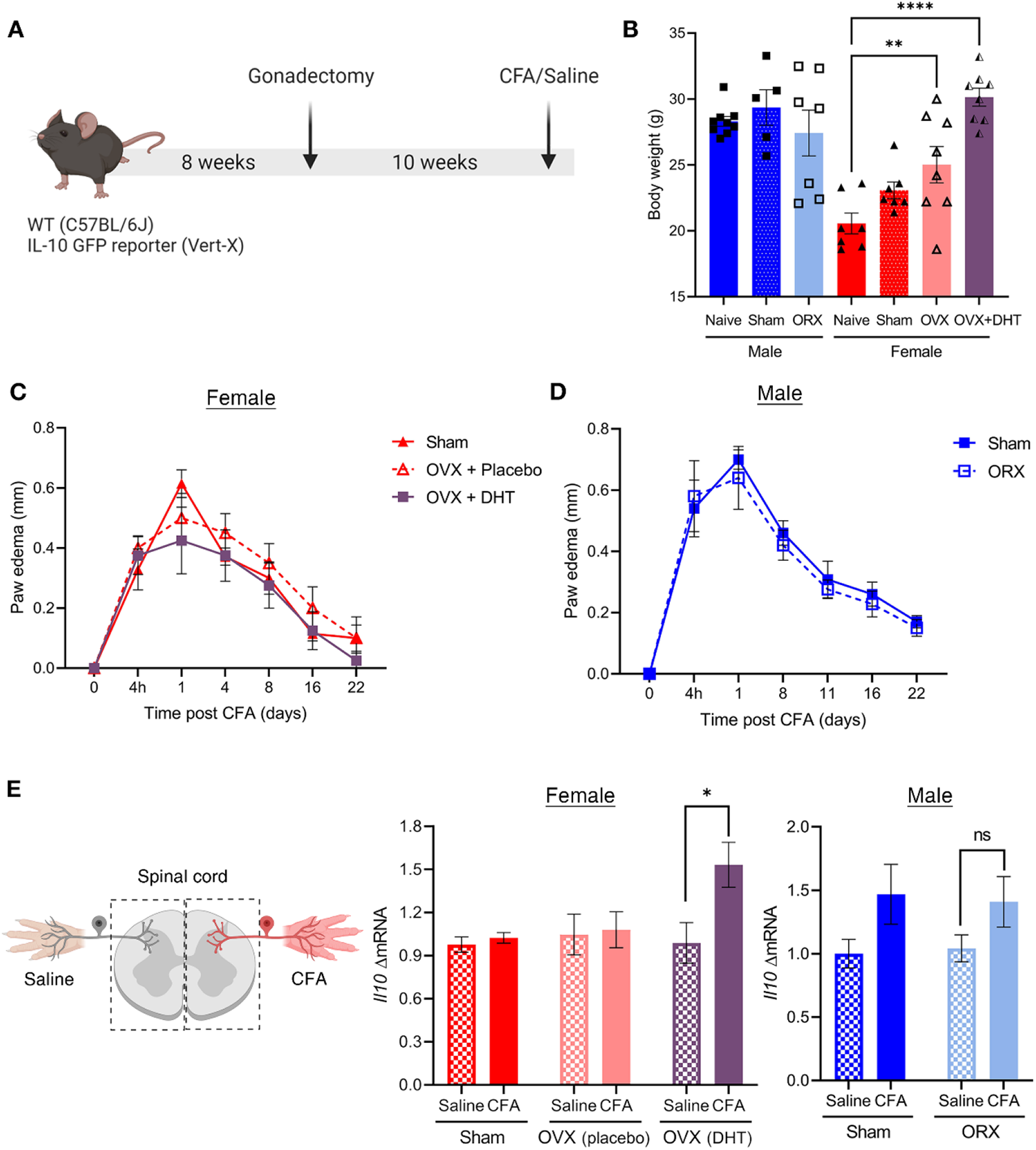
Sex hormones do not regulate paw edema resolution and spinal *Il10* expression. (A) Brief description of the timeline of experiments. (**B**) Body weights of sex hormone-manipulated mice (10 weeks after surgery). (**C, D**) The thickness of paw edema after CFA treatment. Female: sham (n=7), OVX+PBO (n=4), OVX+DHT (n=4). Male: sham (n=5), ORX (n=5). (**E**) Relative *Il10* mRNA level in the lumbar spinal cord of sex hormone-manipulated WT mice, 7 days after Saline or CFA treatment. Female: Sham (n=7), OVX + Placebo (n=8), OVX + DHT (n=8). Male: Sham (n=5), ORX (n=5). Statistical analysis: Ordinary one-way ANOVA, Tukey’s multiple comparisons (**B**), Repeated measures two-way ANOVA, Tukey’s multiple comparisons (**C, D**), two-way ANOVA, Tukey’s multiple comparison s (**E**). *p < 0.05, **p < 0.01, ****p < 0.001, ns, not significant. Symbols represent means and error bars represent SEM (**C, D**). Error bars show mean and SEM (**E**). Abbreviation: OVX, ovariectomy; PBO, placebo; DHT, dihydrotestosterone; ORX, orchiectomy.

### Supplementary Tables

**Supplemental Table 1.**
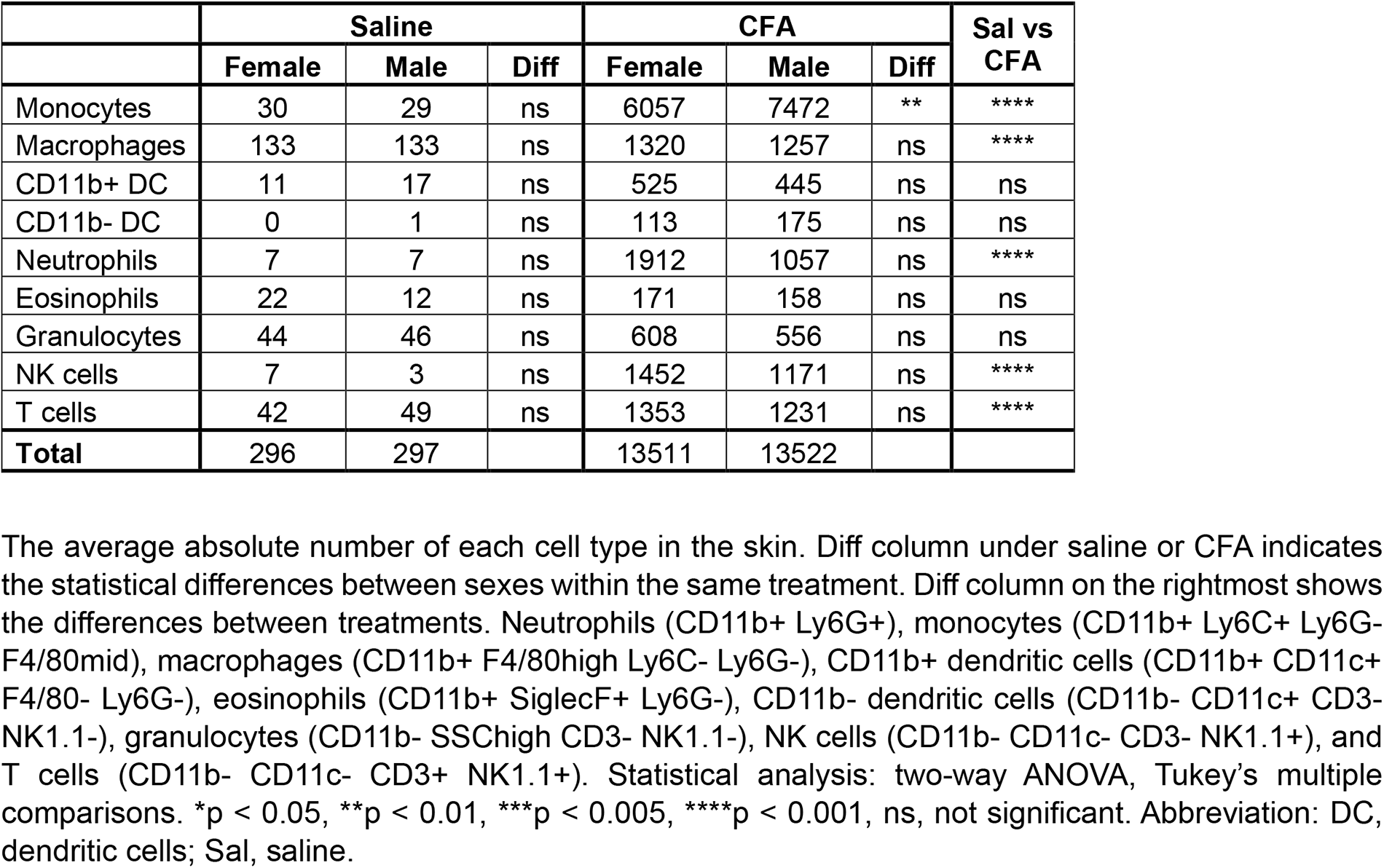
The absolute number of immune cells in the plantar skin.

**Supplemental Table 2.** Percentage of GFP positive immune cells in the CFA-treated skin.

| GFP+ (IL-10-producing) | % of immune cells |  |  |
| --- | --- | --- | --- |
|  | Female | Male | Diff |
| Monocytes | 2.53 | 5.16 | **** |
| Macrophages | 0.99 | 1.17 | ns |
| CD11b+ DC | 0.05 | 0.08 | ns |
| CD11b- DC | 0.00 | 0.01 | ns |
| Neutrophils | 0.01 | 0.01 | ns |
| Eosinophils | 0.00 | 0.02 | ns |
| Granulocytes | 0.22 | 0.25 | ns |
| NK cells | 0.05 | 0.07 | ns |
| T cells | 0.44 | 0.43 | ns |
| **Total** | 4.29 | 7.20 | **** |

Percentage of each GFP+ cell population in total skin immune cells in CFA-treated skin. Diff column indicates the statistical differences between sexes. n=3 (each group). Statistical analysis: two-way ANOVA, Tukey's multiple comparison. \*\*\*\*p < 0.001, ns, not significant. Abbreviation: DC, dendritic cells.

**Supplemental Table 3.** Baseline characteristics of study participants (n=245) from the AURORA study.

| Characteristic | All | Women | Men |
| --- | --- | --- | --- |
| Participants, n | 245 | 172 | 73 |
| Age, years, mean (SD) | 39.2 (14.2) | 40.0 (13.8) | 37.5 (15.0) |
| Self-identified race/ethnicity, n (%) |  |  |  |
| Black | 158 (64.5) | 114 (66.3) | 44 (60.3) |
| White | 80 (32.7) | 52 (30.2) | 28 (38.4) |
| Other | 7 (2.8) | 6 (3.5) | 1 (1.4) |
| Education, n (%) |  |  |  |
| HS or less | 82 (33.4%) | 49 (28.5%) | 33 (45.2%) |
| Some college | 106 (43.3%) | 76 (44.2%) | 30 (41.1%) |
| College or higher | 57 (23.3%) | 47 (27.3%) | 10 (13.7%) |
| ADI, mean (SD) | 69.1 (26.4) | 69.0 (25.9) | 69.4 (27.6) |
| IL-10 (pg/ml), mean (SD) | 0.349 (0.328) | 0.322 (0.329) | 0.413 (0.318) |
| Baseline pain severity<sup>a</sup>, mean (SD) | 6.8 (2.2) | 7.0 (2.2) | 6.5 (2.2) |
| Previous traumatic events<sup>b</sup>, mean (SD) | 14.5 (7.7) | 15.4 (7.6) | 12.3 (7.4) |

<sup>a</sup>Baseline pain severity was measured using the numeric rating scale (0-10 NRS).
<sup>b</sup>Previous traumatic events prior to the currently assessed trauma exposure were assessed via the life events checklist. Abbreviations: SD, standard deviation; HS, high school; ADI, area deprivation index.

**Supplemental Table 4.** Regression analyses assessing the relationship between peritraumatic circulating IL-10 levels and pain severity three months following trauma exposure in multiethnic women and men (n=245).

| | $\beta$ | S.E. | p value |
| --- | --- | --- | --- |
| Intercept | -0.458 | 1.957 | 0.815 |
| Age | 0.031 | 0.014 | 0.031 |
| Sex | -0.744 | 0.456 | 0.104 |
| IL-10 | -0.455 | 0.210 | 0.031 |
| ADI | 0.003 | 0.009 | 0.773 |
| Baseline pain | 0.372 | 0.093 | <0.001 |

Pain severity was measured via a 0-10 numeric rating scale (NRS). Participants' self-identified race/ethnicity, the study site that the participant was enrolled at, and the assay plate for IL-10 processing (to adjust for potential batch effects) were also included as categorical variables in the model, but for simplicity (due to multiple categories), were not included in the table. IL-10 levels were log2 transformed before including in the model. ADI = area deprivation index, S.E. = standard error.

**Supplemental Table 5.** Regression analyses in women and men assessing the relationship between peritraumatic circulating IL-10 levels and pain severity three months following trauma exposure in multiethnic women and men (n=245).

|  | Women (n=172) |  |  | Men (n=73) |  |  |
| --- | --- | --- | --- | --- | --- | --- |
| | $\beta$ | S.E. | p value | $\beta$ | S.E. | p value |
| Intercept | -1.182 | 2.141 | 0.582 | -1.871 | 2.742 | 0.498 |
| Age | 0.032 | 0.017 | 0.066 | 0.037 | 0.028 | 0.193 |
| IL-10 | -0.337 | 0.258 | 0.194 | -0.666 | 0.419 | 0.118 |
| ADI | 0.006 | 0.011 | 0.541 | -0.016 | 0.018 | 0.358 |
| Baseline pain | 0.312 | 0.112 | 0.006 | 0.656 | 0.219 | 0.004 |

Pain severity was measured via a 0-10 numeric rating scale (NRS). Participants' education level was also included as a categorical variable in the model, but for simplicity (due to multiple categories), was not included in the table.

**Supplemental Table 6.**
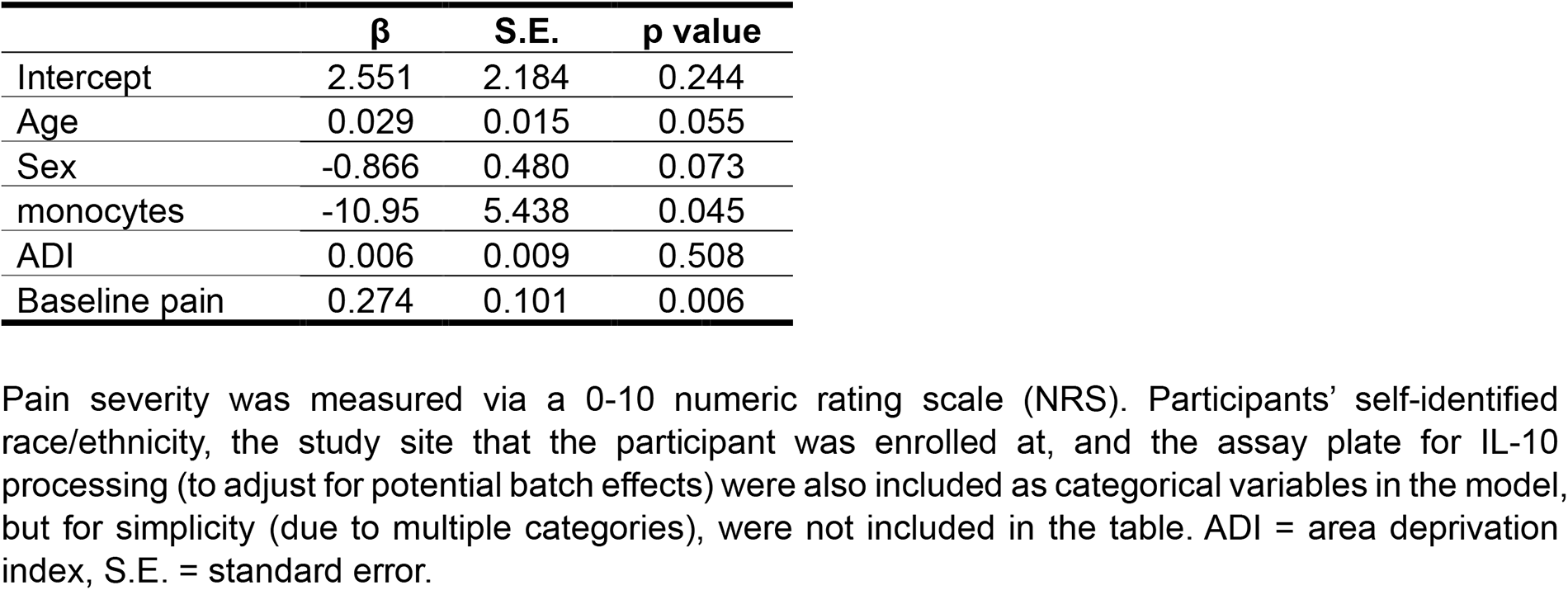
Regression analyses assessing the relationship between peritraumatic circulating monocyte levels and pain severity three months following trauma exposure in multiethnic women and men (n=245).

**Supplemental Table 7.** Percentage of immune cells in the plantar skin of sex hormone manipulated mice.

|  | Female |  |  |  |  |  | Male |  |  |  |
| --- | --- | --- | --- | --- | --- | --- | --- | --- | --- | --- |
|  | Sham |  | OVX+PBO |  | OVX+DHT |  | Sham |  | ORX |  |
|  | Sal | CFA | Sal | CFA | Sal | CFA | Sal | CFA | Sal | CFA |
| Monocytes | 0.14 | 7.31 | 0.17 | 10.80 | 0.20 | 13.57 | 0.12 | 13.08 | 0.07 | 10.36 |
| Macrophages | 4.47 | 4.34 | 2.69 | 6.62 | 2.98 | 3.13 | 1.95 | 3.88 | 2.04 | 5.31 |
| CD11b+ DC | 0.22 | 2.05 | 0.17 | 2.17 | 0.34 | 1.36 | 0.23 | 1.61 | 0.40 | 1.43 |
| CD11b- DC | 0.06 | 0.80 | 0.07 | 1.09 | 0.06 | 0.53 | 0.03 | 0.78 | 0.05 | 0.43 |
| Neutrophils | 0.08 | 6.56 | 0.07 | 2.27 | 0.09 | 4.15 | 0.06 | 3.33 | 0.27 | 2.19 |
| Eosinophils | 0.08 | 0.48 | 0.03 | 0.25 | 0.00 | 0.06 | 0.01 | 0.08 | 0.02 | 0.13 |
| Granulocytes | 0.39 | 4.21 | 0.17 | 2.00 | 0.04 | 2.49 | 0.19 | 2.17 | 0.13 | 3.77 |
| NK cells | 0.10 | 3.36 | 0.03 | 2.41 | 0.22 | 2.07 | 0.33 | 2.34 | 0.39 | 1.68 |
| T cells | 0.40 | 3.39 | 0.39 | 2.48 | 0.48 | 4.15 | 0.36 | 3.33 | 0.58 | 2.19 |
| **Total** | 5.94 | 32.50 | 3.80 | 30.10 | 4.41 | 31.50 | 3.27 | 30.60 | 3.95 | 27.50 |

Each cell type in total live cells %
Percentage of each cell type within skin live cells of sex hormone-manipulated mice. The number of cells were quantified using flow cytometry. Three mice were pooled into one sample for flow cytometry experiments. N=1 or every group. Abbreviation: OVX, ovariectomy; DHT, dihydrotestosterone; ORX, orchiectomy; Sal, Saline; PBO, placebo; DC, dendritic cells.

**Supplemental Table 8.** Percentage of GFP positive immune cells in the CFA-treated skin of sex hormone manipulated mice.

| GFP+ | Female |  |  | Male |  |
| --- | --- | --- | --- | --- | --- |
|  | Sham | OVX+PBO | OVX+DHT | Sham | ORX |
| Monocytes | 0.98 | 2.21 | 6.35 | 5.69 | 4.88 |
| Macrophages | 0.83 | 1.75 | 2.00 | 1.91 | 2.88 |
| CD11b+ DC | 0.27 | 0.30 | 0.15 | 0.23 | 0.24 |
| CD11b- DC | 0.05 | 0.05 | 0.03 | 0.06 | 0.00 |
| Neutrophils | 0.00 | 0.00 | 0.03 | 0.03 | 0.04 |
| Eosinophils | 0.05 | 0.04 | 0.00 | 0.00 | 0.00 |
| Granulocytes | 0.22 | 0.57 | 0.98 | 0.92 | 0.47 |
| NK cells | 0.14 | 0.17 | 0.06 | 0.12 | 0.08 |
| T cells | 0.55 | 0.36 | 1.08 | 0.73 | 0.61 |
| **Total** | 3.09 | 5.46 | 10.68 | 9.71 | 9.20 |

% of live immune cells in skin
Percentage of each cell type within skin live cells of sex hormone-manipulated mice. The number of cells were quantified using flow cytometry. Three mice were pooled into one sample for flow cytometry experiments. N=1 or every group. Abbreviation: OVX, ovariectomy; DHT, dihydrotestosterone; ORX, orchiectomy; Sal, Saline; PBO, placebo; DC, dendritic cells.

**Supplemental Table 9.** Reagent table.

| **Reagent** | **Source** | **Identifier** | **Location** |
| --- | --- | --- | --- |
| **Antibodies** |  |  |  |
| LIVE/DEAD™ Fixable Blue Dead Cell Stain Kit | Invitrogen™ | L34962 | Carlsbad, CA |
| anti-CD45, BUV563 | BD Bioscience | 612924 | Franklin Lakes, NJ |
| anti-F4/80, BV750 | BD Bioscience | 747295 | Franklin Lakes, NJ |
| anti-CD11b, APC-Fire750 | BioLegend | 101261 | San Diego, CA |
| anti-Ly6G, BV711 | BD Bioscience | 563979 | Franklin Lakes, NJ |
| anti-CD11c, PE-Cy7 | BioLegend | 117317 | San Diego, CA |
| anti-CD19, BV785 | BioLegend | 115543 | San Diego, CA |
| anti-NK1.1, BUV805 | BD Bioscience | 741926 | Franklin Lakes, NJ |
| anti-CD3, APC-Fire810 | BioLegend | 100267 | San Diego, CA |
| anti-Ly6C, BV421 | BioLegend | 128031 | San Diego, CA |
| anti-FcεR1, Super bright 600 | Invitrogen™ | 63-589-882 | Carlsbad, CA |
| anti-SiglecF, Alexa fluor 647 | BioLegend | 155519 | San Diego, CA |
| anti-CD11c, BV650 | BioLegend | 117339 | San Diego, CA |
| Anti-Interleukin-10 antibody produced in goat | Millipore Sigma | I5145 | St. Louis, MO |
| IL-10Rα Antibody (E-6) | Santa Cruz | 28371 | Dallas, Texas |
| Recombinant Anti-NeuN antibody | abcam | 177487 | Cambridge, UK |
| Goat anti-Mouse IgG (H+L) Secondary Antibody, Alexa Fluor™ 488 | Invitrogen™ | A-11001 | Carlsbad, CA |
| Goat anti-Rabbit IgG (H+L) Secondary Antibody, Alexa Fluor™ 594 | Invitrogen™ | A-11001 | Carlsbad, CA |
| anti-PGP9.5 | Neuromics | GP14014 | Edina, MN |
| Goat anti-Guinea Pig IgG (H+L) Secondary Antibody, Alexa Fluor™ 568 | Invitrogen™ | A-11075 | Carlsbad, CA |
| **Chemicals and Buffers** |  |  |  |
| Deoxyribonuclease I from bovine pancreas | Millipore Sigma | DN25 | St. Louis, MO |
| Collagenase D | Millipore Sigma | 11088858001 | St. Louis, MO |
| RPMI-1640 Medium | Millipore Sigma | R7509 | St. Louis, MO |
| Gibco™ HBSS (10X), no calcium, no magnesium, no phenol red | Thermo Fisher | 14-185-052 | Waltham, MA |
| Brilliant staining Buffer | BD Bioscience | 563794 | Franklin Lakes, NJ |
| CD16/CD32 Rat anti-Mouse, Unlabeled, Clone: 2.4G2, BD | BD Bioscience | 553142 | Franklin Lakes, NJ |
| UltraComp eBeads™ Compensation Beads | Invitrogen™ | 01-2222-41 | Carlsbad, CA |
| Freund's Adjuvant, Complete | Sigma Aldrich | F5881 | St. Louis, MO |
| Placebo | Innovative Research of America | C-111 | Sarasota, FL |
| 5α-DIHYDROTESTOSTERONE | Innovative Research of America | A-161 | Sarasota, FL |
| cOmplete™, Mini, EDTA-free Protease Inhibitor Cocktail | Millipore Sigma | 11836170001 | St. Louis, MO |
| Fast SYBR™ Green Master Mix | Applied Biosystems | 4385612 | Waltham, MA |
| Clodronate liposome | Liposoma | C-005 | Amsterdam, The Netherlands |
| **Kits** |  |  |  |
| cDNA using high-capacity cDNA Reverse Transcription kit | Applied Biosystems | 4368814 | Waltham, MA |

### Supplementary methods

#### AURORA study: Human participants

All human participants included in the current analyses were participants of the AURORA study, a large longitudinal study of adverse neuropsychiatric outcome development following trauma exposure^2^. Enrollment for this study took place between September 2017 and June 2021. Participant enrollment occurred at 23 urban emergency departments across the US, in the immediate aftermath of traumatic collision exposure, when the individual presented for care. Enrollment focused on individuals ages 18-75 presenting within 72 hours of trauma exposure (predominately motor vehicle collision trauma) who could speak and read English, were oriented to time-space and able to follow the enrollment protocol, were physically able to use a smartphone and possessed a smartphone for >1 year. We excluded participants with a solid organ injury Grade >1 based on American Association for the Surgery of Trauma criteria, significant hemorrhage, need for a chest tube or operation with general anesthesia, or likely to be admitted for >72 hours. All 245 participants included in the current analyses met these criteria, provided informed consent, completed baseline assessments in the emergency department, and provided blood samples for plasma and RNA collection.

#### Human plasma collection

Blood samples were collected in the early aftermath of trauma exposure. Blood was collected into EDTA tubes and immediately centrifuged, and plasma samples were aliquoted and frozen at -20°C to -80°C at the enrollment site. All samples were batch shipped to the University of North Carolina at Chapel Hill Bio-specimen Processing facility on dry ice and stored at -80°C until use.

#### MSD Analysis of Cytokines

Human plasma cytokine (IL-10) was measured in multiplex using Meso Scale Discovery’s V-PLEX Proinflammatory Panel 1 Human chemiluminescence-based assay (MSD, Gaithersburg, MD, USA, catalog # K15049D-1). Plasma samples were thawed on the day of sample processing and diluted with the kit’s sample dilution buffer to 1:2 according to manufacturer recommendations. All assays were performed in duplicate and run in batches of 2 or 3 plates with the same lot number (Lot: Z0047517). In summary, manufacturer-pre-coated plates were washed three times with washing buffer (0.05 % Tween-20 in 1X PBS) before adding samples and standards. Samples for duplicate wells were diluted using 55 µl of the respective plasma in the kit’s sample dilution buffer at a 1:2 ratio. Of this dilution, 25 µl of the standard (including blanks of just the kit’s sample dilution buffer) and plasma samples were added to the assigned wells and incubated for two hours at room temperature on an orbital microtiter plate shaker (IKA MTS 2/4 microtiter shaker Prod. No.: 0003208001). After incubation, plates were washed 3 times, and detection antibodies were added to all the wells and incubated at room temperature for 2 hours. After incubation, the plates were washed 3 times with washing buffer, and 150 µl of reading buffer was added to each well. Plates were immediately analyzed using a QuickPlex SQ 120 instrument (MSD) and the DISCOVERY WORKBENCH^®^ 4.0 software. The intra-and inter-assay coefficient of variation between samples and plates was ≤ 15%. The few samples with higher variation were repeated or excluded from the analysis. Consistent with previously published literature for healthy participants plasma IL-10 levels were detected at an average concentration of 0.349 pg/ml (SD = 0.328 pg/ml).

#### RNA collection, sequencing, and CIBERSORT

To estimate the proportions of immune cell types in each participant, we used RNA sequencing data from the AURORA cohort and a bioinformatic method of identifying cell types based on RNA expression profiles called CIBERSORT^3^. Blood was collected into RNA PAXgene tubes at the same time as EDTA sample collection, in the immediate aftermath of trauma exposure, at the time of enrollment into the AURORA study. After blood collection, RNA PAXgene tubes were incubated at room temperature for 4 hours and then frozen between -20°C and -80°C. Samples were batch shipped to the University of North Carolina at Chapel Hill Bio-specimen Processing facility on dry ice and stored at -80°C until use. Total RNA was isolated using the PAXgene blood miRNA kit (Qiagen, Germantown, MD) and RNA was stored at -80°C until use. RNA concentration and purity were measured using Qubit RNA Assay Kits (ThermoFisher, Waltham, MA) and a 4200 TapeStation System (Agilent Technologies, Santa Clara, CA). The average RNA integrity (RIN) score across the cohort was 7.9, with 92% of samples having a RIN score ≥7. Four samples had RIN values below 4 and were discarded while the remainder were used to prepare RNA sequencing libraries. Template libraries for total RNA sequencing were produced from 600ng total RNA using Genomics Universal Plus Total

RNA Seq Library kits with rRNA and globin depletion (TECAN, Morrisville, NC) according to manufacturer’s specifications. Libraries were multiplexed in groups of 96 and sequenced on a Illumina NovaSeq S4 200 cycle kit (Illumina, San Diego, CA) using paired end reads (100x) at the University of North Carolina at Chapel Hill High Throughput Sequencing Facility. Raw sequencing reads were aligned to the human hg38 genome assembly using STAR (version 2.7.6a)^4^, then quantified using Salmon (1.4.0). Raw read counts were then upper quantile normalized and log2 transformed. The subsequent analysis used only genes that had at least 10 reads in any given library. Outliers were identified via principal component analysis. To verify the appropriate male/female categorization based on genetic attributes, median *XIST* RNA read counts were compared between women and men. CIBERSORT^3^ was adopted to impute gene expression from the transcriptomics data and provide an estimation of the proportion of different immune cell types including B cell subtypes, plasma cells, T cell subtypes, NK cells, monocytes, macrophage subtypes, dendritic cell subtypes, mast cell subtypes, eosinophils, and neutrophils.

#### Animal pain assessment

Mechanical pain sensitivity threshold was measured using von Frey filaments as previously described^5,6^. Briefly, mice were placed in transparent boxes (10 × 10 × 10 cm) with opaque partitions to prevent interaction with other mice. After a 30-minute habituation period, a series of calibrated von Frey filaments were applied perpendicularly to the plantar surface of the hind paw with sufficient force to bend the filaments for 5 seconds. Brisk paw withdrawals or flinching was considered a positive response. In the absence of a response, the filament of the next greater force was applied. The tactile stimulus producing a 50% likelihood of withdrawal was determined using the “up-down” calculating method. Behavioral testing was performed by experimenters blinded to the treatments, genotypes, and sexes. Paw thickness was measured using an electronic caliper. The edema was calculated by the delta of the thickness before and after CFA.

#### Intraplantar injection

Five microliters of 1 mg/ml neutralizing anti-IL-10 (I5145, Sigma, St. Louis, MO) antibody or saline were injected twice on the CFA-treated plantar skin of mice under anesthesia (isoflurane 2.5%) on days 14 and 17 after the CFA treatment. Twenty microliters of clodronate liposome (C005, Liposoma, Amsterdam; The Netherlands) or vehicle (saline) were injected on the CFA-treated skin under anesthesia (isoflurane 2.5%) 14 days after the CFA treatment.

#### Single cell suspensions & Flow cytometry

Mouse skin immune cells were isolated as previously described^7,8^. Mice were euthanized 7 days after saline/CFA injection. Intraplantar skin tissues were collected and placed in the ice-cold 1X Gibco™ HBSS (HBSS (10X), 14-185-052, Waltham, MA) immediately after collection. Two or three paw skins were pooled together to obtain > 20,000 immune cells from the CFA-treated tissues. Tissues were diced into 1 mm-sized pieces to facilitate enzymatic digestion and then transferred to a 15 ml tube filled with the enzyme cocktail, which is constituted of 5 ml of RPMI-1640 Medium (R7509, Sigma, St. Louis, MO), 1 mg/ml DNase I (DN25, Sigma-Aldrich, St. Louis, MO), and 1 mg/ml Collagenase D (11088858001, Sigma, St. Louis, MO). Enzymatic digestion was performed at 37°C for 2 hours. Digested tissues were then minced and filtered through the 70 μm cell strainer by grinding tissue gently with the plunger end of the 10 ml syringe. After centrifugation at 400 g for 10 minutes at 4°C, supernatants were discarded, and the pellets were resuspended with 2 ml of PBS.

For staining of cell surface markers, cells were first stained with 1 μL/mL of LIVE/DEAD Blue (L34962, Invitrogen, Carlsbad, CA) for 15 min on ice. Then, Fc-blocking antibodies CD16/CD32 (553142, BD, Franklin Lakes, NJ) were added to single-cell suspension prior to surface staining. Cells were then stained with the surface marker antibodies (**Supplemental Table 9**) in flow cytometry staining buffer (PBS, 1% BSA) and brilliant staining buffer (563794, BD, Franklin Lakes, NJ). Cells were incubated at 4°C for 1 hour in the dark. Cells were analyzed with a 5 laser (16UV-16V-14B-10YG-8R) Cytek® Aurora – Spectral Flow Cytometry (Cytek Biosciences, Fremont, CA) located in the MSU Flow Cytometry Core Facility, and the data were unmixed with SpectroFlo® software and analyzed using FlowJo™ v10.8 Software (BD Life Sciences). Ultracomp eBeads^TM^ (01-2222-41, Invitrogen, Carlsbad, CA) stained with antibodies were used as the reference controls for unmixing, except for eGFP. The eGFP reference control was prepared using splenocytes from an IL-10-eGFP mouse. Splenocytes from the eGFP mouse were analyzed by Cytek Aurora and only the purified signature of eGFP was exported as FCS file format and later imported as an eGFP reference control. To prevent interference from the autofluorescence, unmixing algorithms in the SpectroFlo software was used to extract autofluorescence from unstained cells.

#### High dimensional data analysis of flow cytometry data

Flow cytometry data were analyzed by using previously established workflow^9^. First, all data were imported into FlowJo, and only Live CD45 positive cells were exported as FCS files. These files were imported to R(v.4.2.0)^10^ and RStudio(2022.02.2+485) for further analysis. Using R/Bioconductor packages, FlowSOM, and CATALYST, immune cells were computationally clustered based on their markers. The resulting data were processed for dimensionality reduction and visual representation with UMAP. Based on heatmaps and UMAP created from the analysis, we manually merged and annotated clusters. Through these steps, immune cell populations in the skin were defined computationally.

#### Gonadectomy and sex hormone treatment

Ovariectomies: C57BL/6J and Il10^GFP^ female mice were anesthetized with isoflurane and ovaries were removed using two separate flank incisions (∼5 mm). For sham surgeries, ovaries were left intact. In hormone replacement studies, either a placebo or dihydroxy-testosterone (DHT) tablet (C-111, A-161, Innovative Research of America, Sarasota, FL) were placed subcutaneously between the scapulae at the time of ovariectomy. Orchiectomies: C57BL/6J and Il10^GFP^ male mice were anesthetized with isoflurane and testis were removed using two separate scrotum incisions. For sham surgeries, testes were left intact. Gonadectomy was performed at 8-10 weeks of age and mice which underwent surgeries were introduced in further experiments at least 10 weeks later.

#### RNA isolation and gene expression assessment

Mice were terminated by CO_2_ and perfused with ice-cold PBS. Skins of the hind paw, spinal cord, and spleen tissues were frozen immediately after the collection and stored at -80⁰C until preparation. Following sonication, RNA was isolated using a modified version of the TRIzol/chloroform extraction method^11^ and dissolved in nuclease free water after incubation in dry bath at 65⁰C. RNA concentration was determined by Qubit 4 Fluorometer (Invitrogen) using the RNA broad range kit (Q10211, Invitrogen). 500ng of RNA was converted into cDNA using a high-capacity cDNA Reverse Transcription kit (4368814, Applied Biosystems, Thermo Fisher Scientific). The expression of *Il10*, *Il10ra*, *Gapdh, Ccr2, Plac8, Ly6g, and Cma1* was assessed using the validated primers (Mm.PT.58.13531087 exon 3-5, Mm.PT.11980819 exon 1-2, Mm.PT.39a.1 exon 2-3, Mm.PT.58.14116710, Mm.PT.58.11720458, Mm.PT.58.30498043 exon 2-3, and Mm.PT.58.45856134 exon 1-2) (Integrated DNA Technologies, Coralville, IA) and SYBRgreen (#4385612, Applied Biosystems, Thermo Fisher Scientific). *Gapdh* was used as a housekeeping gene and values from control samples were normalized to 1.

#### Immunohistochemistry: Dorsal root ganglion

Mice were perfused with ice-cold PBS and 4% paraformaldehyde, and lumbar dorsal root ganglions (DRG) were harvested and placed into 4% paraformaldehyde for fixation and cryoprotected. The tissues were cut into 8 µm thickness sections on a Leica CM3050 S cryostat. As previously described^6^, sections were incubated with mouse monoclonal anti-IL-10Rα (1:100, sc-28371, Santa Cruz, Dallas, Texas) and polyclonal rabbit anti-NeuN (1:500, ab177487, Abcam, Cambridge, UK) overnight at 4⁰C. Subsequently, the sections were incubated with Alexa fluor 488 goat anti-mouse (1:500, A-11029, Invitrogen, Carlsbad, CA) and Alexa fluor 594 goat anti-rabbit (1:500, A-11037, Invitrogen, Carlsbad, CA). Omitting primary antibodies resulted in the absence of staining. Sections were visualized on confocal microscopy (CTR400 Leica).

#### Immunohistochemistry: Skin intra-epidermal fiber staining

Plantar surfaces of the hind paws were collected and fixed in 2% paraformaldehyde at 4⁰C for 24 hours. Then the tissue was transferred to 20% sucrose in PBS and then incubated in 30% sucrose in PBS for at least 7 days. Once the tissues sink to the bottom of the solution, tissues were frozen in OCT:30% sucrose (2:1) and kept at -80°C. They were sliced (20 µm) with a cryostat and immediately mounted on Superfrost Plus slides (Fisher Scientific). Slides were dried and stored at -80°C until use. The slide sections were rehydrated with PBS and tissues were permeabilized with 0.3% Triton X-100 in PBS under RT for 15 minutes. Then, they were blocked with blocking buffer (5% bovine serum albumin, 3% normal goat serum, 0.3% Triton X-100 in PBS) under RT for 1 hour. Tissues were stained with guinea pig anti-mouse PGP9.5 (Neuromics, GP14104, 1:300) and mouse anti-mouse IL-10Rα (Santa Cruz, sc-28371, 1:100) diluted in PBST and incubated overnight at 4°C. In next day, after washing, the slides were incubated with host-matched secondary antibody goat anti-mouse Alexa Fluor 488 (Invitrogen, A11001, 1:300) (Ex: 499 nm; Em: 520 nm) and goat anti-guinea pig Alexa Fluor 568 (Invitrogen, A11075, 1:300) (Ex: 579 nm; Em: 603 nm) under dark, overnight at 4°C. Sections were washed twice with PBS and deionized water and coverslipped with ProLong Gold (ThermoFisher).

Images were acquired with Nikon C2 Upright confocal microscope. Two lasers (488 nm and 560nm) were used to excite the fluorochromes and 505LP, 560LP, 525/50, and 600/50 are the emission filters used. Images were captured with Nikon PlanApo 20x/0.75NA Objective without immersion medium. To prevent crossover errors, sequential imaging was selected. Z series was performed with a step size of 1 μm along the 15 μm-thick scanning area. After the acquisition, every stack image was processed as a Maximum Intensity Projection Image. Adobe Photoshop or Fiji/Image J (Fiji Is Just ImageJ) was applied to image direction and magnification adjustment, along with a small brightness and cell count correction.

